# Apigenin as a Candidate Prenatal Treatment for Trisomy 21: Effects in Human Amniocytes and the Ts1Cje Mouse Model

**DOI:** 10.1101/495283

**Authors:** Faycal Guedj, Jeroen LA Pennings, Ashley E Siegel, Fatimah Alsebaa, Lauren J Massingham, Umadevi Tantravahi, Diana W Bianchi

## Abstract

Human fetuses with trisomy 21 (T21) have atypical brain development that is apparent sonographically in the second trimester. Prenatal diagnosis provides a potential opportunity to begin treatment *in utero*. We hypothesize that by analyzing and integrating dysregulated gene expression and pathways common to humans with DS and mouse models we can discover novel targets for therapy. Here, we tested the safety and efficacy of apigenin (4’, 5, 7-trihydroxyflavone), identified using this approach, in both human amniocytes from fetuses with T21 and in the Ts1Cje mouse model. The experiments compared treated to untreated results in T21 and euploid cells, as well as in Ts1Cje mice and their wild-type littermate controls. T21 cells cultured with apigenin (2µM) had significantly reduced oxidative stress and improved antioxidant defense response *in vitro*. Apigenin (333-400 mg/kg/day), mixed with chow, was initiated prenatally to the dams and fed to the pups over their lifetimes. There was no significant increase in birth defects or pup deaths resulting from prenatal apigenin treatment. Apigenin significantly improved several developmental milestones and spatial olfactory memory in Ts1Cje neonates. In addition, we noted sex-specific effects on exploratory behavior and long-term hippocampal memory in adult mice, with males showing significantly more improvement than females. Global gene expression analyses demonstrated that apigenin targets similar signaling pathways through common upstream regulators both *in vitro* and *in vivo*. These studies provide proof-of-principle that apigenin has therapeutic effects in preclinical models of Down syndrome.

**ONE SENTENCE SUMMARY:** As a candidate prenatal treatment for Down syndrome, apigenin improved oxidative stress/antioxidant capacity imbalance and reduced pathways associated with inflammation in human cells while improving aspects of behavior in the Ts1Cje mouse model.

## INTRODUCTION

Screening for trisomy 21 (T21) is universally offered as part of routine obstetric care in most developed countries. With the implementation of cell-free DNA sequencing of maternal plasma, the positive predictive values are on the order of 80% in the general obstetric population and ∼92% in the high-risk population (1). In continuing pregnancies, knowledge that the future child will have Down syndrome (DS) may affect the parents’ choice of where to deliver, and provide opportunities for both family education and to meet with pediatric subspecialists before the child’s birth (2). For the past six years our laboratory has suggested consideration of prenatal diagnosis as a potential opportunity to treat the fetus *in utero* (3,4). This concept, however, has many unique challenges. Treatment cannot harm the pregnant woman or her fetus, and any therapeutic agent must cross both the placental and blood-brain barriers and improve postnatal outcomes in the baby.

Prenatal sonographic and post-mortem studies demonstrate that atypical brain growth is first detectable in second trimester fetuses with T21, resulting in significant reduction of neurogenesis, synaptogenesis, axonal growth and myelination (5-9). One study has shown that during the third trimester, fetuses with T21 have atypical patterns of habituation to a repeated auditory stimulus, suggesting that functional and sensory deficits are present prior to birth (10). We hypothesize that prenatal treatment given to the pregnant woman as soon as a diagnosis of T21 is made will result in more typical fetal brain growth and development.

Until recently, almost all preclinical and clinical trials in mouse models and people with DS have been conducted in adolescents and adults due to safety concerns. Three main strategies have been used: neural stem cell transplantation, pharmacotherapy and environmental enrichment combined with physical exercise (11). To date, 11 molecules have been tested with little evidence of success in humans with DS (11-13). Potential reasons for this failure may be related to the fact that these therapeutic interventions were carried out too late, and not during the prenatal and early postnatal critical periods for brain development (14,15). To date, no prenatal treatment studies have been reported in pregnant women carrying fetuses with T21. A limited number of prenatal treatment studies using fluoxetine, maternal choline supplementation and the neuroprotective peptides NAP and SAL have been described using the Ts65Dn mouse model of DS (16-18).

In our previous studies, we integrated gene expression data from nine different cellular and tissue sources in both humans with DS and mouse models to identify common dysregulated signaling pathways and cellular processes (19,20). We demonstrated that pathway abnormalities associated with Down syndrome were the result of gene-dosage specific effects and the consequence of a global stress response with activation of compensatory mechanisms (20). To counteract these genome-wide abnormalities, we used the Connectivity Map database (21) (www.broadinstitute.org/CMap) to discover molecules that could be repurposed to rescue the transcriptome and promote more typical brain development in individuals with DS (20). One of the molecules that had the most consistent negative scores (hence, negating the dysregulated gene expression signatures in DS) across tissues and species was apigenin (4’, 5, 7-trihydroxyflavone).

Apigenin is a molecule of interest because it has no known toxicities. It is a naturally occurring compound that is present in chamomile flowers, parsley, celery, peppermint, and citrus fruits (22,23). In animal studies, apigenin has been shown to cross the blood-brain barrier. It has potent anti-oxidant, anti-inflammatory, and anti-apoptotic properties (23,24). In murine microglia that have been activated by interferon gamma, apigenin decreased the levels of IL-6 and TNF-alpha via its effect on phosphorylation of STAT1 (25). This is of particular interest because both humans with DS and mouse models show consistent evidence of interferon abnormalities (26,27). In a double transgenic mouse model for amyloid precursor protein and presenillin 1 proteins, oral intake of apigenin for three months resulted in reduction of fibrillar amyloid deposits and improvement in learning and memory deficits (28).

Here we performed a proof of principle study to test the potential prenatal therapeutic effects of apigenin *in vitro* on human amniocytes and *in vivo* on one model of DS, the Ts1Cje mouse. Although we have recently described the results of an extensive comparison of three major mouse models of DS (26), the experiments reported here were initiated before the comparative study reported in Aziz *et al*. was completed. Whereas Ts1Cje mice are more mildly affected than Ts65Dn mice, we did not select the latter model because affected males are sterile. This requires the trisomic chromosome to be passed through an affected mother, altering the intrauterine environment in which the fetus develops. This can confound the postnatal evaluation of pup development. Furthermore, Ts65Dn mice harbor a large segmental trisomy of non-orthologous human *Hsa21* genes from mouse *Mmu17* for which the impact on the phenotype is still unclear.

## RESULTS

### Effects of Apigenin Treatment on Human Amniocytes

#### Optimal Dose Selection

Separate analyses of apigenin effects on euploid and T21 amniocytes showed similar trends. High doses of apigenin (4 and 5 µM) negatively impacted cell proliferation, with ∼10% reduction in total cell number in euploid amniocytes (U=0 and 6 respectively, p<0.05, Mann-Whitney test) and between 15 to 30% reduction in T21 amniocytes (*p=0.08* at 4µM and *p<0.01* at 5 µM, respectively) (Supplementary Figure 1). Based on these data, we selected doses between 0 and 4 µM for further evaluation of oxidative stress and antioxidant capacity in T21 and euploid amniocytes.

#### Oxidative Stress and Antioxidant Capacity

The average % of DNA in the tail of the untreated T21 amniocytes (14.1±2.6%, Sum of ranks=30, U=0, p=0.016, Mann-Whitney test) was significantly higher than in the euploid amniocytes (5.3±1.1%, Sum of ranks=15) (Figures 1A-B). In treated T21 amniocytes, apigenin significantly reduced the percent of DNA in the tail in a dose-dependent manner (6.7±1.0%, *p=0.06* for 1 µM of apigenin; 5.1±0.8%, *p=0.03* for 2 µM; 2.7±0.8%, *p<0.01* for apigenin 4 µM) (Figures 1A-B). Although not statistically significant, apigenin treatment also reduced the % of DNA in the tail of euploid amniocytes at 2 and 4 µM (2.8±0.8%, *p=0.09* and 2.9±1.2%, *p=0.11* respectively) (Figures 1A-B).

**Figure 1:**
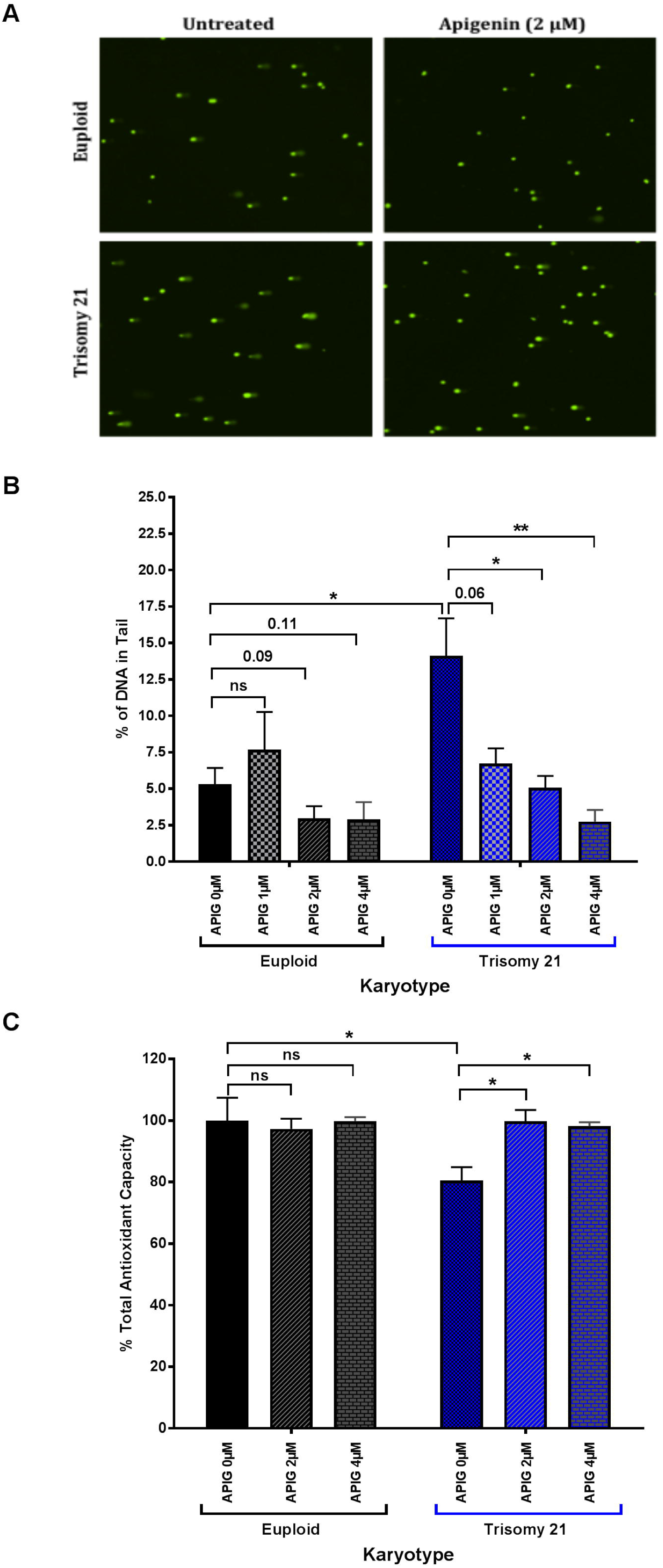
Effects of apigenin treatment on oxidative stress and antioxidant capacity in Trisomy 21 (T21) and euploid amniocytes. **(A)** COMET assay representative images in untreated and apigenin-treated (2 µM) T21 and euploid amniocytes. **(B)** Percent of DNA in tail was analyzed in T21 (N=5) and euploid (N=5) amniocyte cell lines untreated and treated with increasing doses of apigenin (1, 2 and 4 µM). A total of 300-500 cells were analyzed for each cell line and drug dose. Apigenin significantly reduces the % of DNA in the tail in a dose-dependent manner in T21 amniocytes. **(C)** Total Antioxidant Capacity (TAC) measured as absorbance at 490 nm was normalized to 100 % in untreated euploid amniocytes. The percent of TAC in untreated and apigenin-treated (2 and 4 µM) T21 and euploid amniocytes was analyzed. T21 amniocytes exhibit reduced % TAC compared to euploid amniocytes. Apigenin treatment normalized TAC in T21 amniocytes to the level of euploid amniocytes. * (p<0.05), ** (p<0.01).

Two-way ANOVA analysis of the percent (%) of DNA in the tail of both euploid and T21 amniocytes highlighted a significant effect of karyotype (F=4.7, *p=0.04*), treatment (F=7.6, *p<0.001*), as well as treatment x karyotype interaction (F=3.9, *p=0.02*).

Untreated T21 amniocytes also exhibited significantly lower anti-oxidant capacity (80.6±4.2%, *p<0.05*) compared to euploid amniocytes (100.0±7.4%) (Figure 1C). Treatment with 2 and 4 µM of apigenin significantly increased the total anti-oxidant capacity in T21 amniocytes and restored it to levels that were close to euploid amniocytes (99.8±3.7% and 98.1±1.3%, *p>0.05*) (Figure 1C).

#### Global Gene Expression

For gene expression studies, we selected a dose of 2 µM because it did not affect cell proliferation and it rescued the oxidative stress/antioxidant capacity in T21 amniocytes. A total of 14 independent samples were used to generate seven sex and age-matched pairs (Supplementary Table 1). Using paired analysis after elimination of redundant probes, we identified over 500 genes that were differentially expressed (DEX) in T21 amniocytes compared to gestational age and sex-matched euploid amniocytes (Supplementary Table 2A). Fifty of these DEX genes mapped to chromosome 21. Total chromosome 21 gene expression changes in untreated T21 versus euploid amniocytes demonstrated that only a small subset of genes was up-regulated in a gene-dosage dependent fashion (76 genes out of 506 total *Hsa21* genes on the Human Transcriptome HT 2.0 array) (Supplementary Table 2A).

In T21 amniocytes, treatment with 2 µM of apigenin did not have a significant effect on the global gene expression as demonstrated by Principal Component Analysis (PCA) (Figure 2A) or on the expression of the DEX genes (Supplementary Tables 2B-C). Even though apigenin had no effect on T21 DEX genes, a closer look at the lists of marginally expressed (MEX) genes (Supplementary Table 3) identified many candidate genes that were reported in the literature to be direct or indirect targets of apigenin, including *CCL2, IL1A, CYP1B, MMP1, SRPINB2, VDR, DUSP5, AURKB, CCKAR, ITGA11, ARMC4, TENM2, DTL*, CDCA7 and *GRB1* (Table 1), thus warranting a further pathway analysis using the MEX gene lists.

**Table 1:**
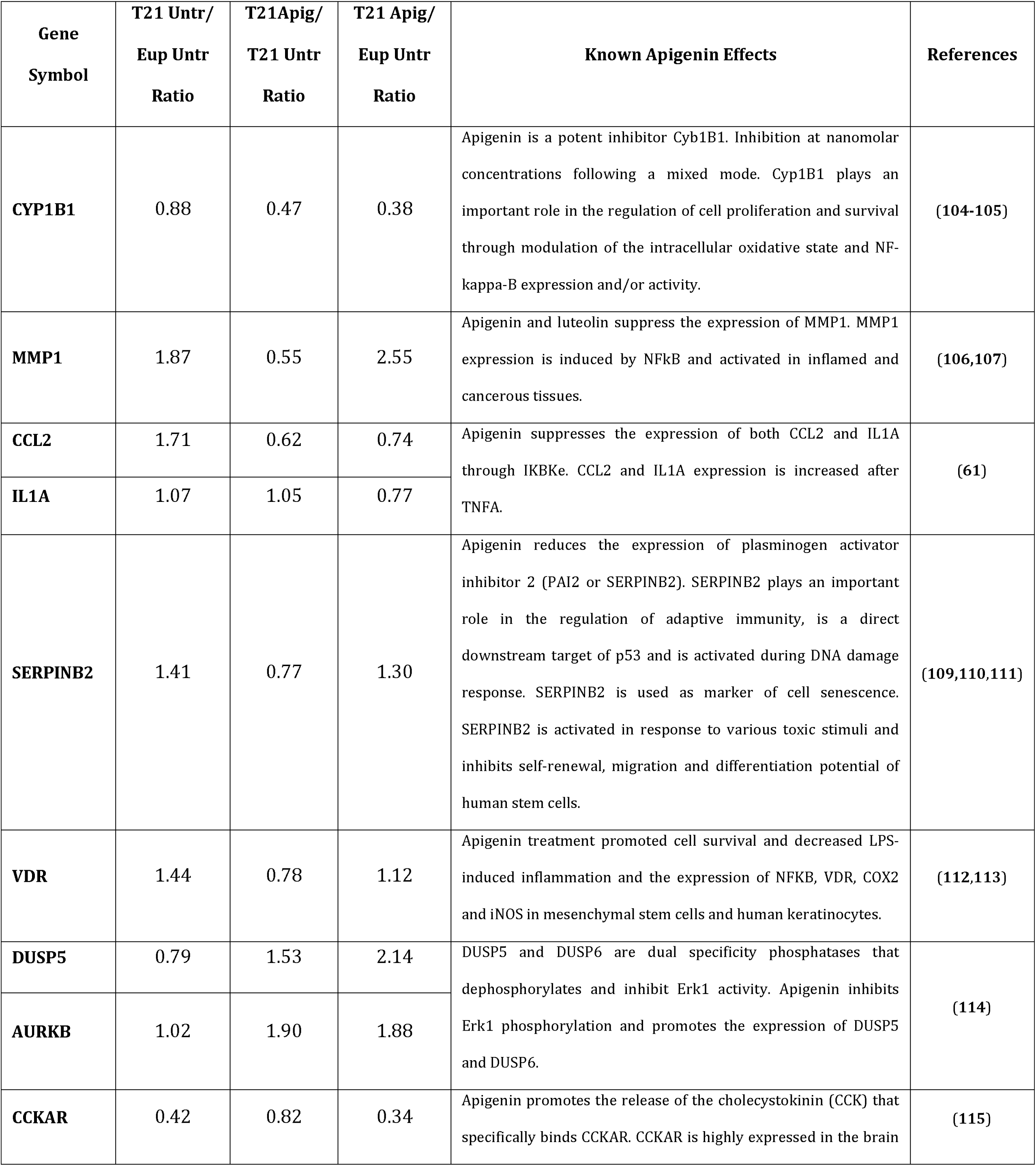
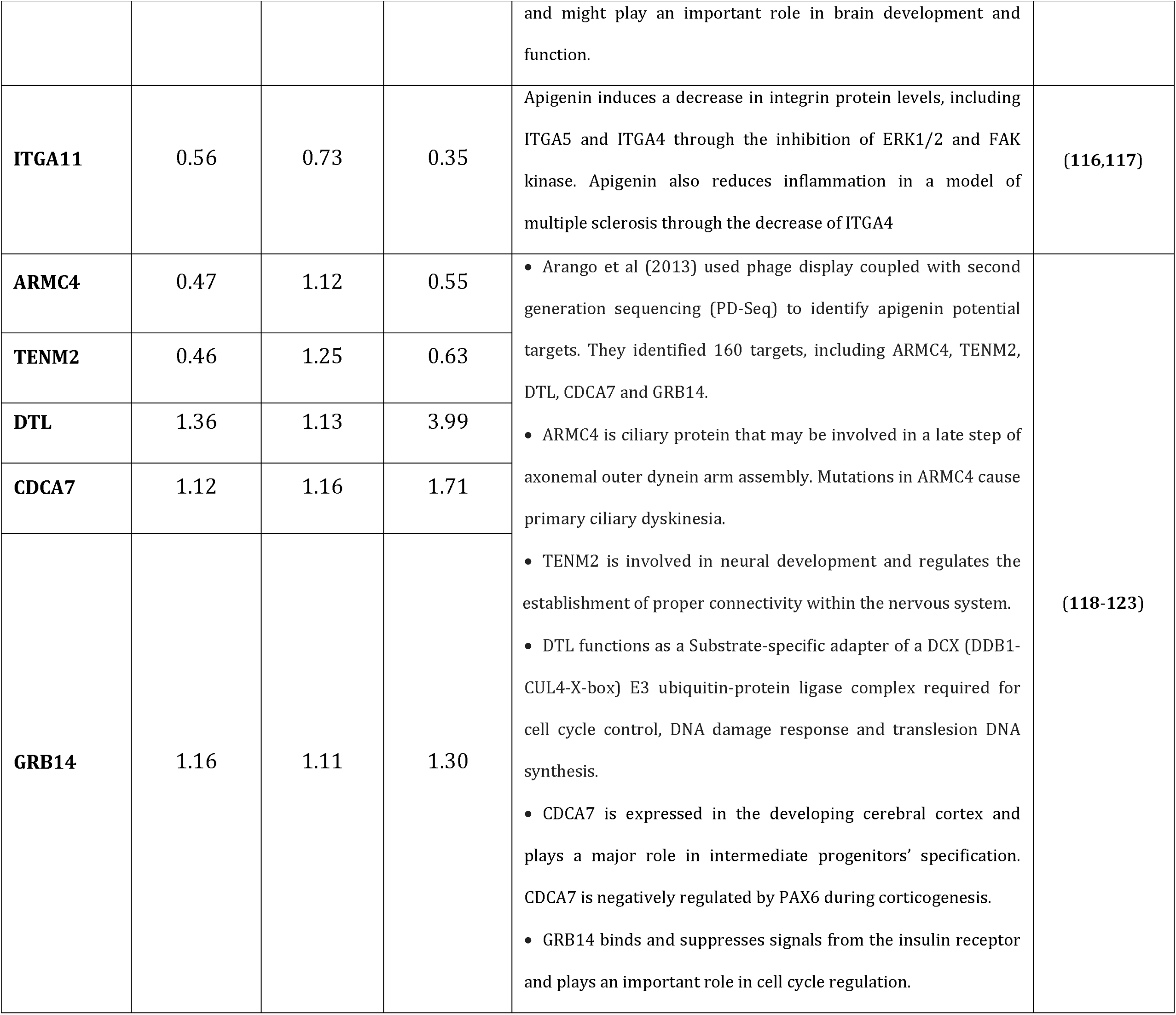
Known Apigenin Targets that are Dysregulated in T21 and Eup Amniocytes After Apigenin Treatment

| Gene Symbol | T21 Untr/<br>Eup Untr<br>Ratio | T21Apig/<br>T21 Untr<br>Ratio | T21 Apig/<br>Eup Untr<br>Ratio | Known Apigenin Effects | References |
| --- | --- | --- | --- | --- | --- |
| **CYP1B1** | 0.88 | 0.47 | 0.38 | Apigenin is a potent inhibitor Cyb1B1. Inhibition at nanomolar concentrations following a mixed mode. Cyp1B1 plays an important role in the regulation of cell proliferation and survival through modulation of the intracellular oxidative state and NF-kappa-B expression and/or activity. | (104-105) |
| **MMP1** | 1.87 | 0.55 | 2.55 | Apigenin and luteolin suppress the expression of MMP1. MMP1 expression is induced by NFkB and activated in inflamed and cancerous tissues. | (106,107) |
| **CCL2** | 1.71 | 0.62 | 0.74 | Apigenin suppresses the expression of both CCL2 and IL1A through IKBKe. CCL2 and IL1A expression is increased after TNFA. | (61) |
| **IL1A** | 1.07 | 1.05 | 0.77 |  |  |
| **SERPINB2** | 1.41 | 0.77 | 1.30 | Apigenin reduces the expression of plasminogen activator inhibitor 2 (PAI2 or SERPINB2). SERPINB2 plays an important role in the regulation of adaptive immunity, is a direct downstream target of p53 and is activated during DNA damage response. SERPINB2 is used as marker of cell senescence. SERPINB2 is activated in response to various toxic stimuli and inhibits self-renewal, migration and differentiation potential of human stem cells. | (109,110,111) |
| **VDR** | 1.44 | 0.78 | 1.12 | Apigenin treatment promoted cell survival and decreased LPS-induced inflammation and the expression of NFkB, VDR, COX2 and iNOS in mesenchymal stem cells and human keratinocytes. | (112,113) |
| **DUSP5** | 0.79 | 1.53 | 2.14 | DUSP5 and DUSP6 are dual specificity phosphatases that dephosphorylates and inhibit Erk1 activity. Apigenin inhibits Erk1 phosphorylation and promotes the expression of DUSP5 and DUSP6. | (114) |
| **AURKB** | 1.02 | 1.90 | 1.88 |  |  |
| **CCKAR** | 0.42 | 0.82 | 0.34 | Apigenin promotes the release of the cholecystokinin (CCK) that specifically binds CCKAR. CCKAR is highly expressed in the brain and might play an important role in brain development and function. | (115) |
| **ITGA11** | 0.56 | 0.73 | 0.35 | Apigenin induces a decrease in integrin protein levels, including ITGA5 and ITGA4 through the inhibition of ERK1/2 and FAK kinase. Apigenin also reduces inflammation in a model of multiple sclerosis through the decrease of ITGA4 | (116,117) |
| **ARMC4** | 0.47 | 1.12 | 0.55 | • Arango et al (2013) used phage display coupled with second generation sequencing (PD-Seq) to identify apigenin potential targets. They identified 160 targets, including ARMC4, TENM2, DTL, CDCA7 and GRB14.<br>• ARMC4 is ciliary protein that may be involved in a late step of axonemal outer dynein arm assembly. Mutations in ARMC4 cause primary ciliary dyskinesia.<br>• TENM2 is involved in neural development and regulates the establishment of proper connectivity within the nervous system.<br>• DTL functions as a Substrate-specific adapter of a DCX (DDB1-CUL4-X-box) E3 ubiquitin-protein ligase complex required for cell cycle control, DNA damage response and translesion DNA synthesis.<br>• CDCA7 is expressed in the developing cerebral cortex and plays a major role in intermediate progenitors' specification. CDCA7 is negatively regulated by PAX6 during corticogenesis.<br>• GRB14 binds and suppresses signals from the insulin receptor and plays an important role in cell cycle regulation. | (118-123) |
| **TENM2** | 0.46 | 1.25 | 0.63 |  |  |
| **DTL** | 1.36 | 1.13 | 3.99 |  |  |
| **CDCA7** | 1.12 | 1.16 | 1.71 |  |  |
| **GRB14** | 1.16 | 1.11 | 1.30 |  |  |

**Figure 2:**
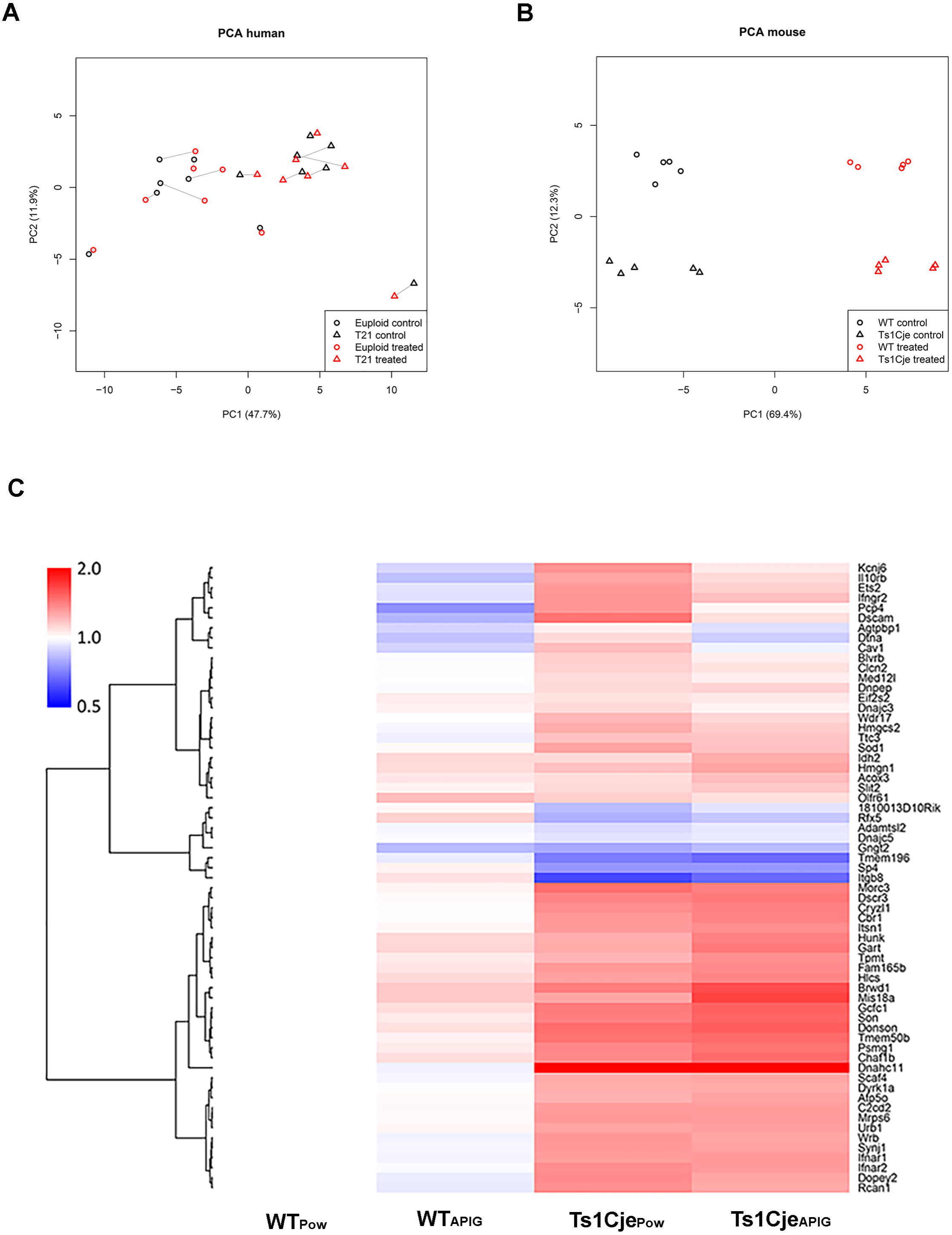
Effects of apigenin treatment on global gene expression *in vitro* and *in vivo*. **(A)** Principal Component Analysis (PCA) of DEX gene expression in untreated and apigenin-treated T21 (N=7) and sex and age matched euploid amniocytes (N=7). Apigenin treatment induced subtle gene expression changes in T21 and euploid amniocytes. Solid lines link each untreated cell line to its treated counterpart. **(B)** PCA of DEX gene expression in untreated and apigenin-treated Ts1Cje (N=5) and Wild-Type (N=5) E15.5 embryonic forebrain. Apigenin induced significant changes in gene expression in both Ts1Cje and WT embryos. These changes seem to be independent of the genotype. **(C)** Heat map demonstrating the effect of apigenin on differentially expressed (DEX) genes in Ts1Cje versus WT embryos. WT_Pow_: untreated WT, Ts1Cje_Pow_: untreated Ts1Cje, WT_APIG_: apigenin-treated WT, Ts1Cje_APIG_: apigenin-treated Ts1Cje.

#### Pathway Dysregulation

Compared to untreated euploid amniocytes, T21 amniocytes showed a positive enrichment of gene sets associated with nucleosome assembly, G1/S mitotic cell cycle transition, RNA Polymerase I-dependent transcription, G-protein signaling, proteolysis, immune response and JAK-STAT signaling. In contrast, negative enrichment of gene sets associated with RNA Polymerase II-dependent transcription, translation initiation, G2/M mitotic cell cycle transition, NOTCH signaling and response to hypoxia was observed (Table 2, Supplementary Tables 4-6).

Apigenin treatment up-regulated RNA polymerase II-dependent transcription and down-regulated the pro-inflammatory response and JAK-STAT signaling in T21 amniocytes to the level of the euploid untreated amniocytes (Table 2). Additionally, apigenin treatment induced an over-compensation of gene sets associated with G2/M cell cycle transition and positive regulation of cell proliferation particularly through Polo-Like kinase pathway; and down-regulation of G-protein signaling when compared to untreated euploid and T21 amniocytes. Apigenin also inhibited NFκB signaling but did not affect other dysregulated pathways in untreated T21 amniocytes (Table 2, Supplementary Tables 4-6).

**Table 2:**
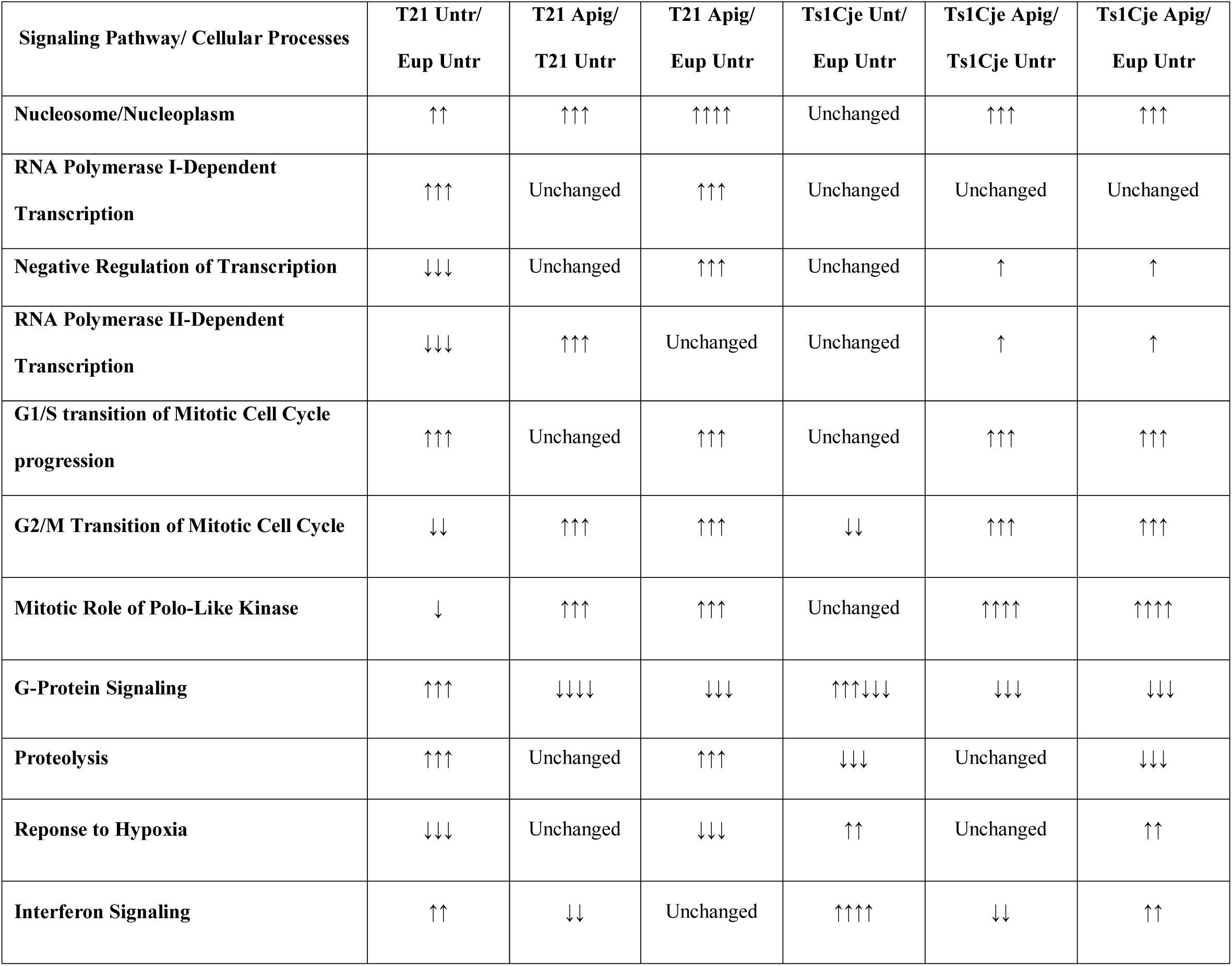
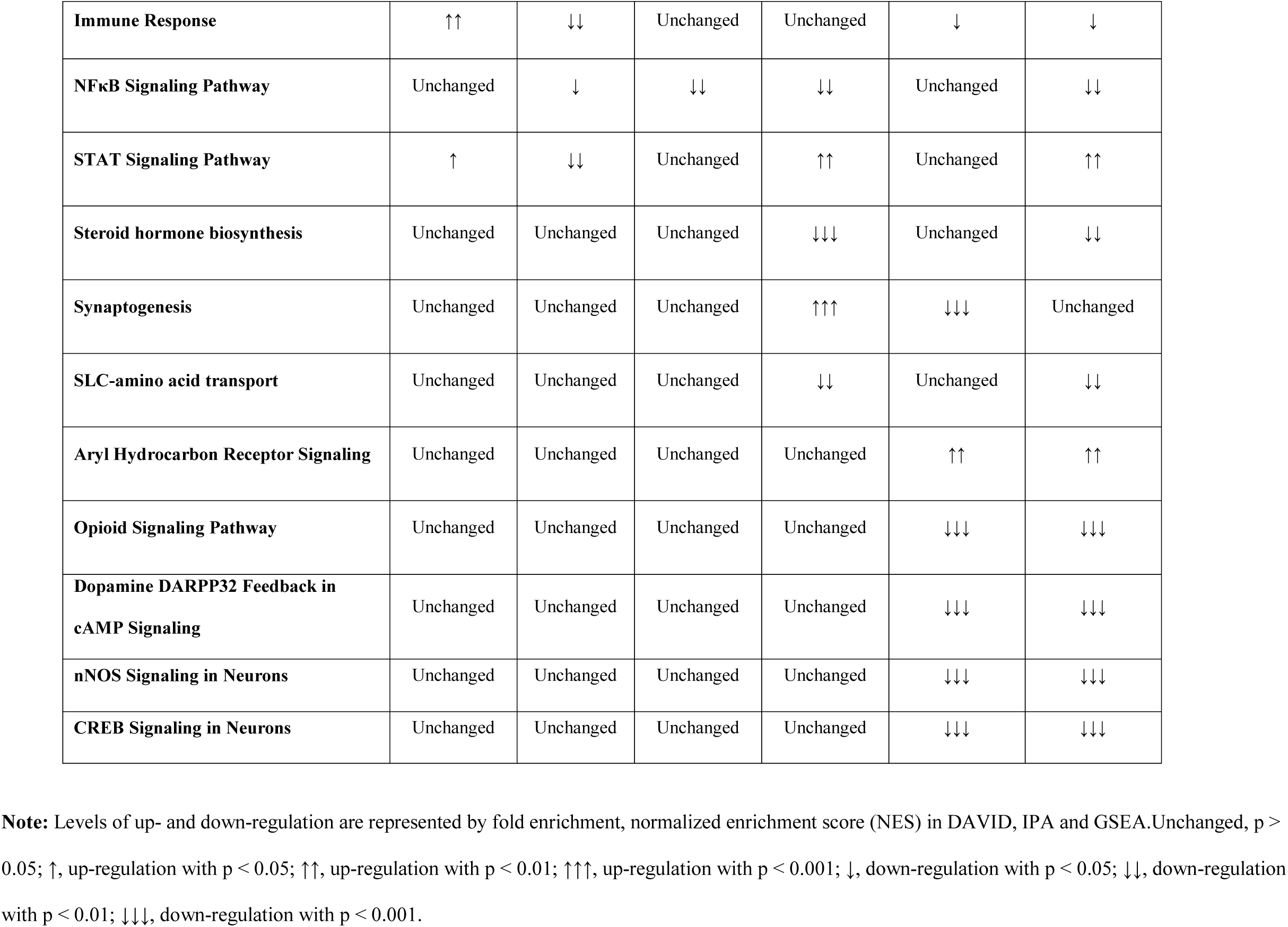
Summary of Dysregulated Signaling Pathways and Cellular Processes in Untreated and Apigenin-treated T21 Amniocytes

#### Upstream Regulators in Untreated and Apigenin-Treated T21 Amniocytes

IPA upstream regulator analysis in untreated T21 amniocytes predicted the activation of VEGFA (chemokine activity and regulation of angiogenesis), APP (neurite outgrowth), interferon gamma (interferon signaling), IKBKB and TLR4 (NFκB signaling). TP53 (DNA damage-repair), PTGER 2 and PTGER4 (inflammation/auto-immunity) and HIF1A (regulation of hypoxia) were predicted to be inhibited (Supplementary Table 7A).

After apigenin treatment, IFNG and IKBKB were predicted to be inhibited in T21 amniocytes to a level closer to untreated euploid cells. Interestingly, HGF (cell proliferation regulation through HGF-Met signaling) and PTGER2, but not PTGER4, were predicted to be the most significantly activated upstream regulator. Additionally, FOXO1 (regulation of homeostasis) and EP400 (cell cycle and transcription regulation) were predicted to be activated to a lesser extent than HGF and PTGER2, followed by E2F1 and E2F2 (transcription activators and G1/S cell cycle transition regulators). In contrast, BNIP3L (response to hypoxia) and IRGM1 (immune response) were predicted to be significantly inhibited (Supplementary Tables 7B-C).

### Effects of Apigenin Treatment in the Ts1Cje Mouse

#### Natural History and Growth

When measured on days E10.5 and E15.5, wild-type (WT) dams fed powdered chow and powdered chow plus apigenin gained similar amounts of weight. At day E15.5, they had an average of 7.2 (17 litters /123 embryos) and 7.8 (8 litters /63 embryos) embryos per litter, respectively.

At E15.5, in the untreated dams, the ratio of WT (49.6%) and Ts1Cje (50.4%) embryos followed Mendelian inheritance. In the apigenin-treated dams, 61.9% of embryos were WT, whereas only 38.1% were Ts1Cje by genotyping (Fisher exact test, *p=0.06*) (Supplementary Figure 2A). These ratios did not change postnatally (Supplementary Figure 2B). Apigenin treatment did not induce any significant effects on the weight or crown-rump length of E15.5 embryonic mice (Supplementary Figures 2C-D). No increase in neonatal deaths or presence of gross congenital anomalies was observed after apigenin treatment in both Ts1Cje and WT pups.

A two-way ANOVA analysis showed a statistically significant effect of the genotype on body weight throughout the pre-weaning period for both untreated and apigenin-treated pups [F(3,1718)=378.7, *p<0.0001*, η^2^=237.6]. Tukey’s multiple comparison test showed that apigenin treatment did not have a significant effect on Ts1Cje pup growth (*p=0.72*) but caused a slight weight increase (*p=0.04*) without significantly affecting body length in WT_Apig_ versus WT_Pow_ (Supplementary Figures 2E-F). Similar to what has previously been observed for body weight, Ts1Cje untreated and apigenin-treated pups were smaller in length compared to their WT littermates.

#### Differentially Expressed (DEX) Genes

For all of the analyses, three groups were compared: 1) the pups of untreated dams compared by genotype (Ts1Cje_Pow_vs. WT_Pow_); 2) Ts1Cje pups that were treated (_Apig_) vs. untreated (_Pow_); and 3) Ts1Cje treated pups vs. untreated WT. In some experiments, WT_Apig_ and WT_Pow_ were also analyzed.

Overall, apigenin treatment induced similar expression changes in both coding and non-coding genes in Ts1Cje_Apig_ (1399 DEXs versus Ts1Cje_Pow_) and WT_Apig_ (906 DEXs versus WT_Pow_). Even though the number of DEX genes was lower in the WT_Apig_, PCA analysis demonstrated that the regulation direction of genes affected by apigenin was similar between Ts1Cje_Apig_ and WT_Apig_ littermates (Figure 2B, Supplementary Table 8C). Importantly, apigenin-treated Ts1Cje embryonic forebrain showed a significant up-regulation of genes implicated in neural stem cell proliferation (*Nestin, Sox2, Sox5, Kif4, Prom1, Pax6, Mcm2, Ect2, Nr2l, Gli3* and *Ccnd2*) and proneural genes (*Neurog1, Neurog2, Nhlh1* and *Nhlh2*) implicated in cell fate determination (Supplementary Table 8C).

In addition to the global gene expression changes induced by apigenin treatment in Ts1Cje embryonic forebrain, we sought to determine if apigenin specifically rescues the expression of DEX genes in Ts1Cje embryos. In Ts1Cje_Pow_ vs. WT_Pow_ embryonic forebrain, 63 protein-coding genes were differentially regulated. Forty-two of these genes mapped to the Ts1Cje aneuploid regions (38 genes on *Mmu16* and 4 on *Mmu12*) (Supplementary Table 8A). Nineteen of the DEXs mapped to chromosomes other than *Hsa21* orthologs. In Ts1Cje_Apig_ vs. Ts1Cje_Pow_ embryonic forebrain (Figure 2C, Supplementary Table 8A-B) the expression of seven genes (*Dscam, Kcnj6, Pcp4, Ets2, Il10rb, Cav1* and *Dtna*) was partially corrected (expression decreased after treatment by >30%) and four genes (*Brwd1, Mis18a, Gart* and *Hunk*) were amplified (expression increased after treatment by ≥30%). These partially corrected and amplified genes were not specifically affected in Ts1Cje_Apig_, but followed similar regulation direction in the WT_Apig_ versus WT_Pow_ (Supplementary Table 8C).

#### Embryonic Forebrain Signaling Pathway Abnormalities

Because of the low number of DEX genes in the Ts1Cje_Pow_/WT_Pow_ comparison, we examined the MEX gene lists (Supplementary Table 9) for pathway analysis using the GSEA, DAVID and IPA databases (Supplementary Tables 10-12).

Similar to what we have described previously (26,29), untreated Ts1Cje embryos exhibited a significant dysregulation of G-protein-coupled receptor activity (mainly olfactory receptor activity) and down-regulation of gene sets associated with cell proliferation (G2/M cell cycle transition), proteolysis, SLC-mediated amino acid transport, NFκB signaling, regulation of translation and steroid hormone synthesis. Additionally, gene sets related to immune response, interferon signaling, Jak-Stat signaling, regulation of hypoxia through Hif1a and synaptogenesis were significantly up-regulated (Table 2, Supplementary Tables 10-12).

Apigenin treatment reduced the immune response in Ts1Cje_Apig_ forebrain and had a pleiotropic effect resulting in a partial improvement of some of the above-mentioned pathways, including Jak-Stat signaling, NFκB signaling and Slc-amino acid transport (Table 2, Supplementary Tables 10-12). Apigenin also induced a significant down-regulation of GPCR signaling but did not have any significant effects on regulation of hypoxia through *Hif1a* (Table 2).

Additionally, gene sets associated with cell cycle progression (G2/M cell cycle transition and Polo-like kinase signaling), DNA-damage repair and kinetochore organization were over-compensated by apigenin treatment. Finally, apigenin significantly inhibited the p53 signaling pathway, opioid signaling, nNOS signaling and Creb signaling pathways (Table 2, Supplementary Tables 10-12).

#### Embryonic Forebrain Upstream Regulators

IPA upstream regulator analysis revealed that untreated Ts1Cje embryonic brains exhibit a significant activation of Ago2 (transcription repression), Tnf (Nfκb signaling), Vegfa (chemokine activity and angiogenesis regulation) and Tp53 (DNA damage-repair). However, Ptger4 was predicted to be inhibited in Ts1Cje_Pow_ versus WT_Pow_ (Supplementary Table 13).

In Ts1Cje embryos, apigenin was predicted to inactivate the action of Tnf and Tp53 but did not affect Ago2 and Vegfa. Apigenin was also predicted to inhibit the action of Bnip3l and Irgm1, and specifically activate Ptger2 (but not Ptger4), Hgf, Ep400, Foxo1, Foxm1 and E2f1, similar to what has been observed in apigenin-treated T21 amniocytes (Supplementary Tables 13-14).

In addition to the upstream regulators that were consistently affected in both T21 amniocytes and Ts1Cje embryonic forebrain after apigenin treatment, several other regulators were predicted to be specifically inhibited by apigenin in the Ts1Cje embryonic forebrain, including Creb1 (activated), Bdnf, Pax6, Gata1 and Il1b (Supplementary Tables 13-14).

#### Neonatal Developmental Milestones

Ts1Cje_Pow_ pups exhibited significant delays versus WT_Pow_ littermates in acquiring early developmental milestones (surface righting, cliff aversion, negative geotaxis and forelimb grasp) and late coordination and sensory maturation milestones (open field, air righting, eye opening and ear twitch) (Table 3, Supplementary Figures 3-4). The percent of Ts1Cje_Pow_ pups reaching criteria (performing the test successfully under 30 s and for two consecutive days) was significantly lower compared to WT_Pow_ littermates (Table 3, Supplementary Figures 3-4).

**Table 3:**
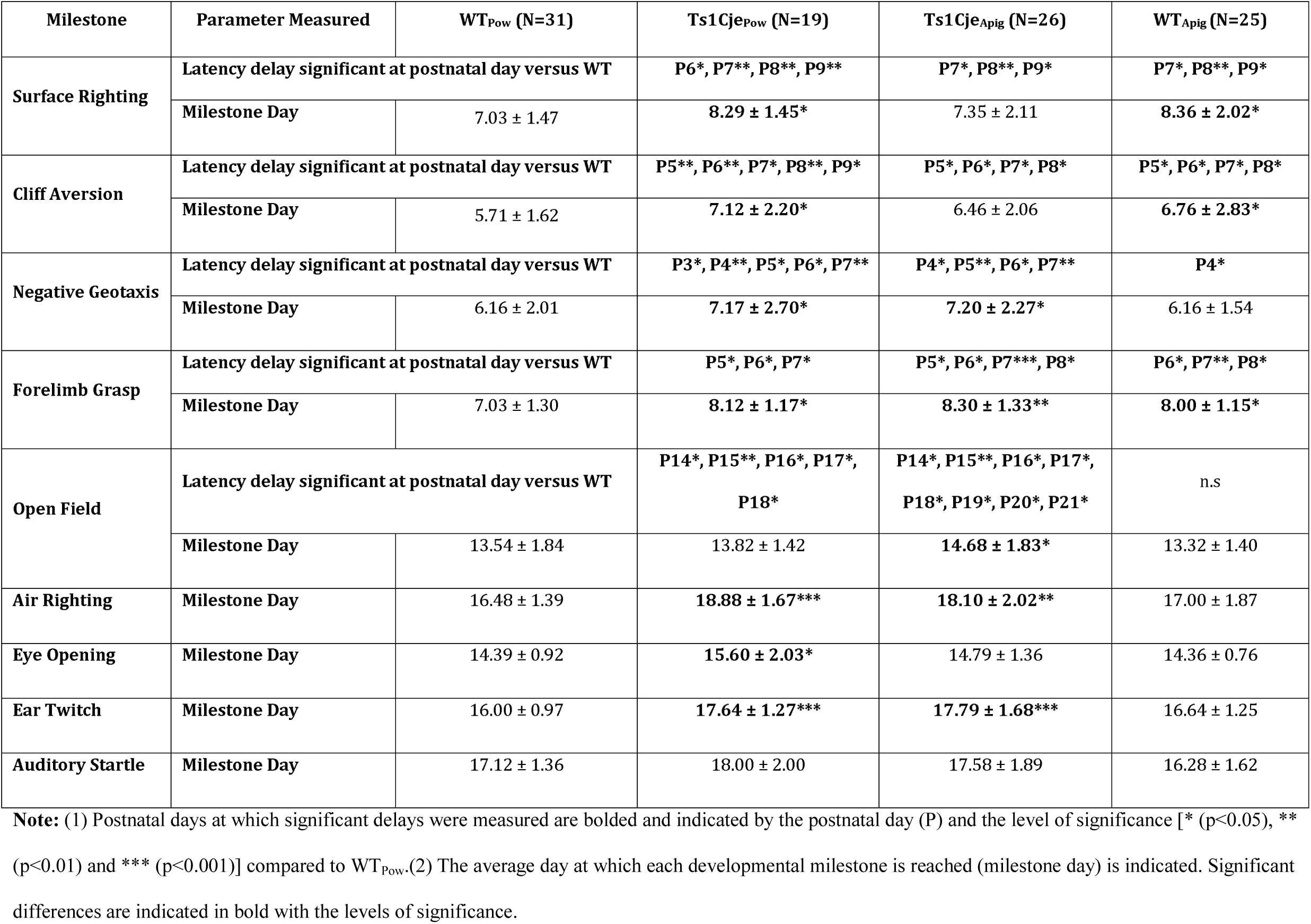
Effects of Apigenin Treatment on Developmental Milestones in Ts1Cje Pups

| Milestone | Parameter Measured | WT<sub>Pow</sub> (N=31) | Ts1Cje<sub>Pow</sub> (N=19) | Ts1Cje<sub>Apig</sub> (N=26) | WT<sub>Apig</sub> (N=25) |
| --- | --- | --- | --- | --- | --- |
| Surface Righting | Latency delay significant at postnatal day versus WT |  | **P6\*, P7\*\*, P8\*\*, P9\*\*** | **P7\*, P8\*\*, P9\*** | **P7\*, P8\*\*, P9\*** |
|  | Milestone Day | 7.03 ± 1.47 | **8.29 ± 1.45\*** | 7.35 ± 2.11 | **8.36 ± 2.02\*** |
| Cliff Aversion | Latency delay significant at postnatal day versus WT |  | **P5\*\*, P6\*\*, P7\*, P8\*\*, P9\*** | **P5\*, P6\*, P7\*, P8\*** | **P5\*, P6\*, P7\*, P8\*** |
|  | Milestone Day | 5.71 ± 1.62 | **7.12 ± 2.20\*** | 6.46 ± 2.06 | **6.76 ± 2.83\*** |
| Negative Geotaxis | Latency delay significant at postnatal day versus WT |  | **P3\*, P4\*\*, P5\*, P6\*, P7\*\*** | **P4\*, P5\*\*, P6\*, P7\*\*** | **P4\*** |
|  | Milestone Day | 6.16 ± 2.01 | **7.17 ± 2.70\*** | **7.20 ± 2.27\*** | 6.16 ± 1.54 |
| Forelimb Grasp | Latency delay significant at postnatal day versus WT |  | **P5\*, P6\*, P7\*** | **P5\*, P6\*, P7\*\*\*, P8\*** | **P6\*, P7\*\*, P8\*** |
|  | Milestone Day | 7.03 ± 1.30 | **8.12 ± 1.17\*** | **8.30 ± 1.33\*\*** | **8.00 ± 1.15\*** |
| Open Field | Latency delay significant at postnatal day versus WT |  | **P14\*, P15\*\*, P16\*, P17\*, P18\*** | **P14\*, P15\*\*, P16\*, P17\*, P18\*, P19\*, P20\*, P21\*** | n.s |
|  | Milestone Day | 13.54 ± 1.84 | 13.82 ± 1.42 | **14.68 ± 1.83\*** | 13.32 ± 1.40 |
| Air Righting | Milestone Day | 16.48 ± 1.39 | **18.88 ± 1.67\*\*\*** | **18.10 ± 2.02\*\*** | 17.00 ± 1.87 |
| Eye Opening | Milestone Day | 14.39 ± 0.92 | **15.60 ± 2.03\*** | 14.79 ± 1.36 | 14.36 ± 0.76 |
| Ear Twitch | Milestone Day | 16.00 ± 0.97 | **17.64 ± 1.27\*\*\*** | **17.79 ± 1.68\*\*\*** | 16.64 ± 1.25 |
| Auditory Startle | Milestone Day | 17.12 ± 1.36 | 18.00 ± 2.00 | 17.58 ± 1.89 | 16.28 ± 1.62 |

**Note:** (1) Postnatal days at which significant delays were measured are bolded and indicated by the postnatal day (P) and the level of significance [\* (p<0.05), \*\* (p<0.01) and \*\*\* (p<0.001)] compared to WT<sub>Pow</sub>. (2) The average day at which each developmental milestone is reached (milestone day) is indicated. Significant differences are indicated in bold with the levels of significance.

Apigenin treatment partially improved several milestones in Ts1Cje pups, including surface righting, cliff aversion, eye opening and air righting. Additionally, apigenin negatively affected strength (forelimb grasp) and motor maturation (open field) but did not affect other milestones (Table 3, Supplementary Figures 3-4).

#### Neonatal Olfactory Spatial Memory

Ts1Cje_Pow_ pups displayed significant olfactory spatial memory deficits manifested by significant increases in the amount of time needed to reach the goal area in trial 1 (166.8±8.2 s) compared to their WT_Pow_ littermates (134.1±11.4 s, *p<0.05*) (Figure 3A). Apigenin treatment significantly reduced the time required for Ts1Cje_Apig_ mice to reach the goal area (139.0±10.3 s) versus Ts1Cje_Pow_ mice (166.8±8.2 s, *p< 0.01*) (Figure 3A). When compared to WT_Pow_ (134.1±11.4 s), Ts1Cje_Apig_ mice exhibited similar performances (139.0±10.3 s, p=0.96) in the homing test. Similarly, in trial 2, Ts1Cje_Pow_ pups also took longer (152.6±9.9 s) to reach the goal area compared to WT_Pow_ pups (113.8±12.3 s, *p=0.02*). Ts1Cje_Apig_ mice found the goal area faster (132.9±13.6 s) than Ts1Cje_Pow_ mice (152.6±9.9 s, *p=0.12*) but this difference did not reach statistical significance (Figure 3B). Additionally, when compared to WT_Pow_ pups (113.8±12.3 s), Ts1Cje_Apig_ mice took slightly longer (132.9±13.6 s s, *p=0.32*) to reach the goal area, but this difference was not statistically significant.

Ts1Cje_Pow_ pups exhibited significant olfactory spatial memory delays in the first and second trials of the homing test compared to their WT_Pow_ littermates. Three Ts1Cje_Pow_ (15.80%) and 11 WT_Pow_ (44%, Chi-square test, *p<0.0001*) pups were able to reach the goal area in trial 1 (Figure 3C). On trial 2, only 6 Ts1Cje_Pow_ pups (31.58%) reached the goal area while 16 WT_Pow_ (64%, Chi-square test, *p<0.001*) were able to successfully perform the test (Figure 3D). Apigenin treatment significantly improved performances in both WT_Apig_ and Ts1Cje_Apig_ in trial 1 (61.9% of WT_Apig_ and 55.6% of Ts1Cje_Apig_, respectively, Chi-square test, *p<0.001*) and trial 2 (85.7% of WT_Apig_ and 50% of Ts1Cje_Apig_, respectively) compared to Ts1Cje_Pow_ (Figures 3C-D). The performance of Ts1Cje_Apig_ neonates was not statistically different from WT_Pow_ pups in trials 1 (*p=0.53*) and 2 (*p=0.07*), respectively.

Two-way ANOVA analysis showed a statistically significant effects of apigenin treatment in both trials 1 [F(3,79)=2.84, *p=0.043*, η^2^=6,561] and 2 [F(3,79)=4.34, *p=0.007*, η^2^=1,2148].

**Figure 3:**
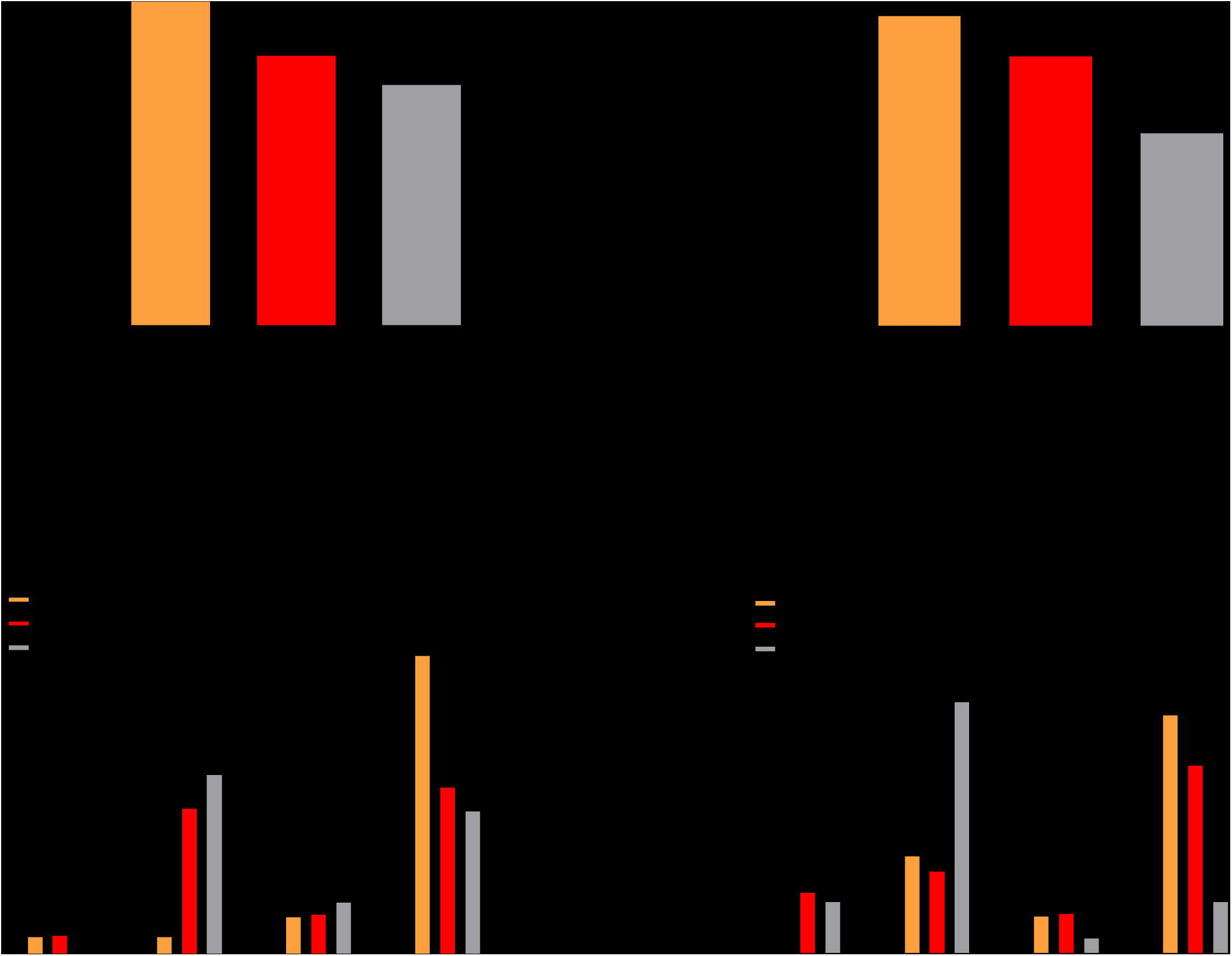
Effects of apigenin treatment on spatial olfactory memory in untreated and apigenin-treated Ts1Cje and WT neonates. The homing test was used to investigate olfactory spatial memory in untreated and apigenin-treated WT and Ts1Cje neonates (WT_Pow_=31, Ts1Cje_Pow_=19, WT_Apig_=25, Ts1Cje_Apig_=26) at postnatal day 12 in two independent trials. **(A-B)** Latency to reach the goal area untreated and apigenin-treated Ts1Cje and WT neonates in trials 1 and 2. Ts1Cje neonates spent significantly more time searching for the goal area compared to their WT littermates suggesting spatial olfactory memory deficits in trisomic mice. Apigenin significantly reduced latency to reach the goal area and improved spatial olfactory memory in Ts1Cje and WT neonates. **(C-D)** Percent of animals reaching the goal area during 60s-time bins in trials 1 and 2. Almost 80% and over 60% of the Ts1Cje neonates and less than 50% of apigenin-treated neonates did not reach the goal area within the 180s trial period in both trials 1 and 2. Apigenin treatment increased the percent of Ts1Cje and WT neonates reaching the goal area during the two first minutes of the trials (0-60 and 61-120 s). * (p<0.05), ** (p<0.01), *** (p<0.001).

#### Adult Exploratory Behavior and Locomotor Activity

Analysis of adult behavior was conducted separately in males and females. In the open field test, the total distance traveled by male Ts1Cje_Pow_ was significantly higher (29,721±1,353 cm) than their WT_Pow_ littermates (23,296±1,019 cm, *p<0.001*) (Figure 4A). Apigenin treatment normalized locomotor behavior in Ts1Cje (distance traveled=25,117±1,443 cm) to the level of WT_Pow_ compared to Ts1Cje_Pow_ animals (*p<0.05*) (Figure 4A).

**Figure 4:**
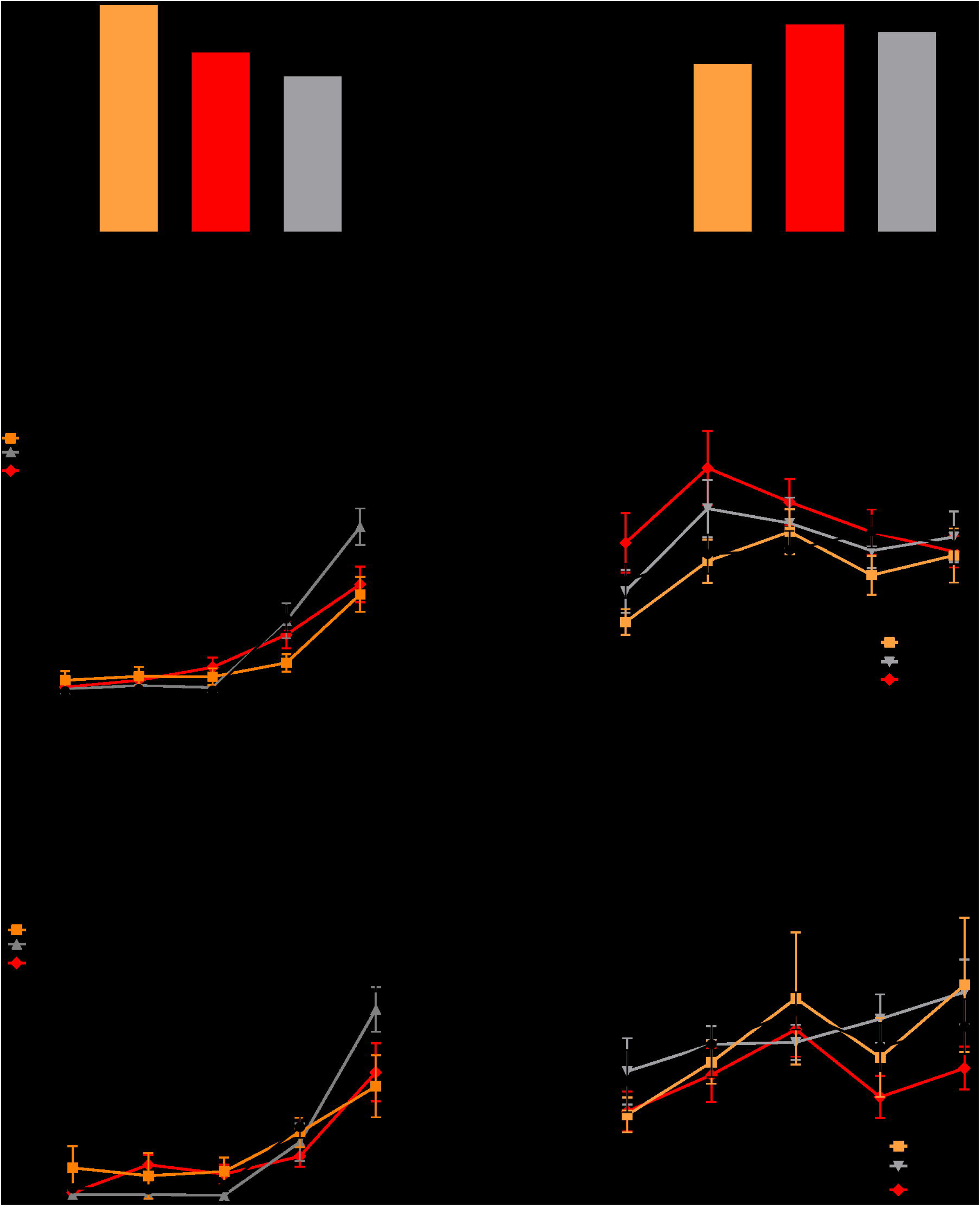
Sex-specific effects of apigenin on exploratory behavior and hippocampal memory in untreated and apigenin-treated adult Ts1Cje and WT males and females. Exploratory behavior and hippocampal-dependent memory were analyzed using the open field and fear conditioning tests, respectively in untreated and apigenin-treated WT and Ts1Cje males (WT_Pow_=13, Ts1Cje_Pow_=12, WT_Apig_=17, Ts1Cje_Apig_=16) and females (WT_Pow_=8, Ts1Cje_Pow_=7, WT_Apig_=14, Ts1Cje_Apig_=12). **(A-B)** Sex specific effects in the open field test. Ts1Cje male but not female mice exhibit hyperactive behavior (higher distance traveled during the 60-min trial) compared to WT littermates. Apigenin treatment rescued exploratory behavior in Ts1Cje male mice but induced hyperactivity in WT and Ts1Cje female mice. **(C-F)** Contextual fear conditioning performances of untreated and apigenin-treated male and female Ts1Cje and WT mice during the training (**C** and **E**) and testing (**D** and **F**) trials. During the testing trial, apigenin treatment significantly increased percent of freezing (improved hippocampal memory) in Ts1Cje males but not in female Ts1Cje during the first two minutes of testing. * (p<0.05), ** (p<0.01), *** (p<0.001).

Ts1Cje_Pow_ females (18,871±1509 cm) did not show alterations in locomotor activity compared to WT_Pow_ (distance traveled=18,133±975 cm) (Figure 4B). Apigenin treatment induced hyperactive behavior in both WT_Apig_ (distance traveled=22,426±1,695 cm) and Ts1Cje_Apig_ females (distance traveled=23,266±1,105 cm, *p<0.01*) (Figure 4B).

#### Adult Hippocampal-Dependent Long-Term Memory

In the fear-conditioning test, during the training phase, Ts1Cje_Pow_ male mice froze significantly less (18.67±3.1%) after receiving the second electrical shock compared to WT_Pow_ (38.40±3.98%, *p<0.001*) (Figure 4C). Similar to their male counterparts, Ts1Cje_Pow_ female mice also showed a significant decrease in % freezing (23.21±6.40%) after receiving the second electrical shock compared to WT_Pow_ (48.10±5.32%, *p=0.013*) (Figure 4E).

In the testing phase, sex differences were observed. Ts1Cje_Pow_ male mice exhibited a lower percent of freezing (21.35±3.03%, *p=0.07*) versus their WT_Pow_ littermates (14.13±2.26%) during the first 60 s, but this difference disappeared during the remaining four minutes of the trial (Figure 4D). Apigenin treatment induced a statistically significant increase in % freezing in the Ts1Cje_Apig_ male mice (28.03±5.23% in the first minute and 41.18±6.54% in the second minute) versus WT_Pow_ and Ts1Cje_Pow_ males during the first two minutes of testing (*p<0.05*) (Figure 4D).

During the testing phase, Ts1Cje_Pow_ females also exhibited a slight non-significant decrease in % freezing (15.89±2.98%) during the first minute compared to WT_Pow_ female mice (21.65±5.30%) (Figure 4F). Apigenin treatment did not affect % freezing in Ts1Cje_Apig_ females (16.44±3.44%) compared to untreated Ts1Cje_Pow_ females (15.89±2.98%) (Figure 4F).

#### Adult Motor Coordination

In the static speed trial of the rotarod, Ts1Cje_Pow_ male mice fell significantly faster at the highest speed (86.40±7.81s at 32 RPM) than WT_Pow_ males (104.30±4.68s at 32 RPM) (Supplementary Figure 5A). However, Ts1Cje_Pow_ females did not show any abnormalities in the rotarod test versus WT_Pow_ littermates (Supplementary Figure 5B). After apigenin treatment, both male and female WT_Apig_ and Ts1Cje_Apig_ fell significantly faster from the rotarod at 32 RPM compared to WT_Pow_ and Ts1Cje_Pow_ untreated mice (*p<0.01*) (Supplementary Figures 5A-B). No negative effects of apigenin were observed at lower speeds (16 and 24 RPM).

In the accelerating speed trial, Ts1Cje_Pow_ male mice also fell significantly faster (224±13.5s) than WT_Pow_ males (268.3±5.46s, *p<0.01*) (Supplementary Figure 5C). A similar trend, although statistically non-significant, was observed in Ts1Cje_Pow_ females (234±19.16s) versus WT_Pow_ females (259.7±11.8s) (Supplementary Figure 5D). Apigenin treatment did not induce any significant adverse effects in both genotypes and genders in the accelerating speed trial (Supplementary Figures 5C-D).

## DISCUSSION

In this study we provide a proof of principle for the safety and efficacy of apigenin both *in vitro* (T21 amniocytes) and *in vivo* (Ts1Cje mouse model). Apigenin reduced oxidative stress and improved anti-oxidant capacity in amniocytes derived from second trimester fetuses with T21. Apigenin also improved some aspects of postnatal behavioral and cognitive outcomes in the Ts1Cje mouse model. Gene expression analyses in both T21 amniocytes and Ts1Cje embryonic forebrain revealed that apigenin consistently promoted the G2/M cell cycle transition (through the up-regulation of Polo-Like kinase signaling) and repressed pro-inflammatory responses and NFκB and G-protein signaling pathways *in vitro* and *in vivo*. This action of apigenin was predicted to be the result of several upstream regulators, particularly PGTER2 and MET receptors, FOXO1, FOXM1, E2F1 and E2F2 transcription factors, and IFNG, TNF, P53 and IKBKB proteins (cell stress and inflammatory response modulators).

### Effects of Apigenin Treatment on Human Amniocytes

#### Improvement in Oxidative Stress/Antioxidant Capacity Imbalance

There is significant transcriptomic, proteomic and biochemical evidence that individuals with DS exhibit a significant imbalance between oxidative stress and physiological response to oxidative stress (i.e. antioxidant capacity) (30-34). This imbalance starts during fetal life, affects multiple organs and might contribute to the atypical brain and cognitive phenotypes in DS (19,35-37). Here, we demonstrated that cultured amniocytes derived from fetuses with DS exhibit an oxidative stress (increased)/anti-oxidant capacity (decreased) imbalance. Apigenin treatment reduced oxidative stress and increased anti-oxidant capacity in T21 amniocytes. Apigenin has been reported to have modulatory effects on oxidative stress and inflammation in different cell models. In cells exposed to reactive oxygen species and external stressors, apigenin has pro-proliferative, anti-inflammatory, anti-oxidant and free radical scavenging effects (38-40).

In healthy cells exposed to internal or external stressors, apigenin plays a cell-specific cytoprotective role by reducing oxidative stress through its direct free radical scavenging action, up-regulation of intracellular anti-oxidant defenses, inhibition of endoplasmic reticulum stress response and activation of MAPK, Nrf2 and UPR (unfolded protein response) signaling cascades (41-43). In cancer cells, apigenin shows anti-proliferative and pro-oxidative stress properties (44-47). Salmani *et al* (48) described apigenin’s mechanisms of action in different types of cancers. In this comprehensive review, it was concluded that apigenin induced cell-type specific molecular changes, but most importantly, apigenin consistently induced cytotoxicity in all cancer cell types studied through increased production of reactive oxygen species and activation of apoptosis.

These data suggest a very complex mechanism of action of apigenin in different cell types through its anti-oxidant, anti-inflammatory, anti-cancer, neuroprotective and anti-neurodegenerative properties (49).

#### Promotion of G2/M Cell Cycle Progression and Suppression of Chronic Inflammation

To get better insights into apigenin’s potential molecular mechanisms of action, we used a transcriptome-based systems biology approach (20). Both *in vitro* and *in vivo*, apigenin treatment improved cell cycle defects (promoting the G2/M cell cycle transition through the Polo-Like kinase cascade), transcriptional dysregulation (promoting RNA polymerase II-dependent transcription), and significantly inhibited the pro-inflammatory response and the NFκB signaling pathway and down-regulated G-protein signaling.

In T21 amniocytes, apigenin induces a significant up-regulation of multiple genes, including *CDK1, NEK2, AURKA, AURKB, CCNB1* and *2, CDC25A* and *C, BIRC5* and *CDC20* that promote entry into M phase (Figure 5A) (50,51). Apigenin inhibits the G2/M transition in malignant cells, repressing proliferation and promoting apoptosis. The regulatory effects on the cell cycle by apigenin in non-malignant cells are poorly studied (52-55).

**Figure 5:**
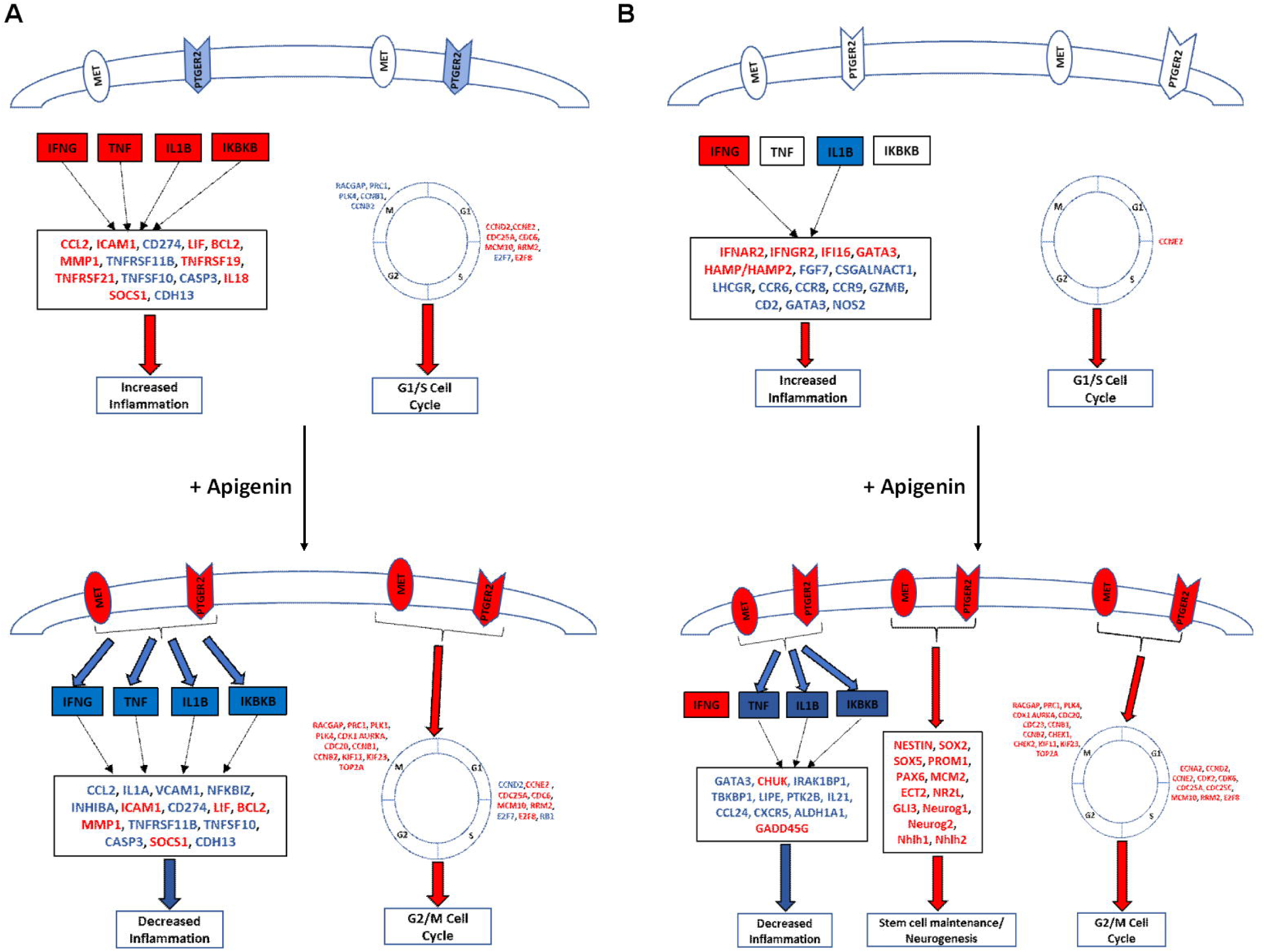
Major Gene Expression Effects of Apigenin Treatment in T21 Amniocytes and the Ts1Cje Mouse Model. (A) Activated and inhibited upstream regulators (predicted by IPA) and genes (in gene expression datasets) in untreated and apigenin treated T21 amniocytes. (B) Activated and inhibited genes and upstream regulators in untreated and apigenin treated Ts1Cje embryonic forebrain. Red=up-regulated, Blue=down-regulated.

Untreated T21 amniocytes exhibit an up-regulation of multiple genes implicated in the pro-inflammatory process and the activation of interferon and NFκB signaling pathways, including *CCL2/MCP1, ICAM1, IL18, MMP1, TNFRSF19, TNFRSF21* (Figure 5A). Although the role of NFκB in the chronic inflammatory response in DS is still poorly studied, several reports have described increased plasma levels of several pro-inflammatory cytokines, including *IL6, IL10, MCP1, TNFA, INFG* and *MMP1* in children and both non-demented and demented adults with DS, suggesting their use as potential biomarkers for disease progression (56-58). Apigenin treatment significantly reduced the expression of several pro-inflammatory genes, including *CCL2, MMP1, IL1A, NFKBIZ, INHIBA* and *VCAM1* in T21 amniocytes (Figure 5A). Multiple *in vitro* studies reported significant reduction of pro-inflammatory molecules (*TNFA, CCL2, IL1A, IL6, IL1B, ICAM1* and *VCAM1*) after apigenin treatment (59-61).

### Effects of Apigenin Treatment in the Ts1Cje Mouse

In this study, we used a high dose of apigenin (>300 mg/kg/day) to evaluate the effects of prenatal and postnatal treatments in the Ts1Cje mouse model of DS. In untreated pregnancies, the ratio of Ts1Cje_Pow_ versus WT_Pow_ embryos followed the expected mendelian distribution. Although not statistically significant, the ratio of Ts1Cje_Apig_ versus WT_Apig_ embryos was slightly lower than the expected mendelian ratio. These results were obtained by examining 123 embryos (17 litters) from untreated dams and only 63 embryos (8 litters) from apigenin fed dams. This observation needs to be confirmed with a larger cohort of embryos, especially since there were no differences in the ratios Ts1Cje_Pow_ versus Ts1Cje_Apig_ neonates.

#### Rescue of the Expression of Several Hsa21 Orthologous Genes in the Fetal Brain

Apigenin treatment induced significant gene expression changes in the embryonic Ts1Cje forebrain. Several *Mmu16* genes were partially compensated by apigenin, including *Dscam, Kcnj6, Pcp4, Ets2, Il10rb, Cav1* and *Dtna*. Although the contribution of some of these genes (i.e. *Il10rb, Cav1* and *Dtna*) to the DS phenotype is still unknown, the remaining genes (*Dscam, Kcnj6, Pcp4* and *Ets2*) have been reported to be highly expressed in the developing brain and overexpressed in brains from individuals with DS. Their overexpression or deletion is associated with cognitive and/or motor deficits in transgenic mouse models (62-67). The molecular mechanism by which apigenin down-regulated these genes is still under investigation.

#### Treatment Targets Similar Pathways in the Ts1Cje fetal brain and T21 amniocytes

Even though the Ts1Cje mouse model did not recapitulate all of the dysregulated pathways described in T21 amniocytes, apigenin treatment targeted similar pathways *in vitro* and *in vivo*. Indeed, apigenin treatment resulted in a significant up-regulation of polo-Like kinase signaling, thus promoting G2/M cell cycle transition, down-regulation of G-protein signaling and reduction of neuro-inflammation through down-regulation of NFκB, Tnf and Il1b signaling (Figure 5B). The anti-inflammatory potency of apigenin has been previously reported in the literature (68-71). Most *in vivo* studies have reported inhibition of the NFκB signaling pathway and suppression of pro-inflammatory cytokines, including Tnfα, Il6 and Ifnγ in different mouse models with different CNS insults, including Parkinson’s and Alzheimer’s diseases. In Ts1Cje mice, apigenin resulted in a significant up-regulation of *Chuk*/*Ikka* (a key kinase negative regulator of NFκB) and down-regulation of *Irak1bp1* and *Tbkbp1* (which plays a role in activating NFκB signaling) (Figure 5B).

Our *in vitro* and *in vivo* gene expression studies suggest that apigenin plays a modulatory role in the cell cycle and has opposite effects in malignant (cell cycle arrest and apoptosis) and non-malignant cells (cell cycle promotion and cytoprotection). More studies are needed to uncover the molecular mechanisms by which these differential modulation mechanisms are activated in cancerous and non-cancerous cells.

#### Treatment is Predicted to Target Similar Upstream Regulators in Ts1Cje Fetal Brain and T21 Amniocytes

Using two independent gene expression datasets (T21 amniocytes and Ts1Cje embryonic forebrain), we could predict that apigenin accomplished its action through similar upstream regulators *in vitro* and *in vivo*, particularly PGTER2 and MET (HGFR) receptors, FOXO1, FOXM1, IFNG, TNF, P53 and IKBKB proteins.

Few studies have investigated the PGE2/PTGER2 and HFG/MET pathways in individuals with DS. Minc-Colomb *et al* (72) reported a reduction in prostaglandin production (PGE2 and PGD2) in fibroblasts from humans with DS and in the kidney and cerebellum of transgenic mice overexpressing *Sod1* gene. In contrast, the levels of COX2 (PGE2/PTGER2 upstream activator) are higher in gingival crevicular fluid from humans with DS and in the cerebral cortex of aging individuals with DS and Alzheimer’s disease (73-75). The role of HGF/MET signaling pathway in the pathophysiology of DS remains unknown.

Most recently, PGE2/PTGER2 and HGF/MET signaling pathways have gained a lot of attention because of their modulatory effects on inflammation, cell stress and cell proliferation. These two pathways act together as a double-edged sword. Depending on the cell type and the downstream targets activated by these two pathways, they can have pro-inflammatory and pro-apoptotic roles or anti-inflammatory, anti-apoptotic and pro-proliferative effects (76,77). Studies in different organ injuries and chronic inflammation models, as well as evidence from knock-out mice for *PTGER2* and *MET*, demonstrated that the activation of PGE2/PTGER2 triggers the HGF/MET signaling to inhibit inflammation and promote cell proliferation and tissue repair (78). The increasing *in vitro* and *in vivo* evidence for the neuroprotective, neurogenic and long-term memory facilitation roles of PGE2/PTGER2 and HGF/MET signaling pathways established them as potential targets for drug development for neuroinflammatory and neurodegenerative conditions, including ischemic brain injury, epilepsy, Alzheimer’s and Parkinson diseases (79-86). In the brain, PTGER2 and MET are expressed in neurons, astrocytes and microglia. However, the mechanism of action by which these receptors are activated and trigger pro-inflammatory/cytotoxic versus anti-inflammatory/neurogenic down-stream effectors is unknown. Based on our gene expression, we hypothesize that apigenin activates HGF/MET signaling (receptor tyrosine kinase) and PGE2/PTGER2 signaling through the G-protein independent pathways, while inhibiting the G-protein dependent pathway.

### Improvement in Several Postnatal Behavioral Outcomes

Prenatal treatment with apigenin improved several aspects of neonatal (surface righting, cliff aversion, air righting and spatial olfactory memory) and adult behavior (open field and fear conditioning). Although the effects of apigenin on early neonatal behavior remained unstudied until now, several studies have demonstrated that apigenin and its most abundant metabolite luteolin, improve behavioral deficits in rodent models of epilepsy and depression. Apigenin and luteolin had significant anxiolytic effects in the elevated plus maze, forced swim test and tail suspension tests (87-90). In other rodent models of neurodegenerative diseases (rotenone-induced Parkinson disease and streptozotocin-induced Alzheimer disease models), epilepsy and cerebral ischemia, apigenin and luteolin significantly improved hippocampal spatial memory in the Morris water maze test (28, 68, 91-95).

Even though treatment with a high dose of apigenin improved exploratory behavior and hippocampal learning, it had negative effects on motor coordination in both Ts1Cje males and females. Anusha *et al* (68) demonstrated beneficial effects of apigenin treatment at low doses (10 and 20 mg/kg, i.p) on motor coordination in a rotenone-induced rat model of Parkinson disease. However, treatment with high doses of apigenin or luteolin inhibited motor function and induced mild sedative effects in mice and rats (89,96). These data suggest the importance of using low to medium doses of apigenin for future preclinical studies in mouse models of DS.

Importantly, prenatal apigenin treatment had sex-specific effects in some adult behavioral paradigms (open field and fear conditioning tests). To our knowledge, no other studies have investigated the sex-specific effects of treatment with polyphenols in rodent models of CNS diseases. Further studies are needed to confirm and investigate the molecular mechanisms of these sex-specific effects. Gradoltto *et al* (97) studied sex and age-specific differences in apigenin metabolism in rats after a single oral 10 mg dose. In mature males, glucuronated and sulfated derivatives of apigenin were present, but in inverse proportions when compared to females and immature males and females, suggesting that sex-specific differences in behavioral outcomes after treatment might be the results of sex-specific differences in metabolism. Our studies stress the importance of evaluating the effects of treatment with polyphenols in *both* males and females because of their estrogenic potential (98).

## STRENGTHS AND WEAKNESSES

Although tremendous advances have been made in understanding the molecular origins of brain and cognitive deficits in DS, human clinical trials conducted so far have been unsuccessful. Some potential reasons for this failure include the methods used to choose the candidate drugs, the use of only the Ts65Dn mouse model to test therapies preclinically and the timing of the interventions. In our previous study, we described a transcriptome-based unbiased approach to identify candidate therapeutic molecules that can rescue the abnormal transcriptome signature in DS (20). Here, we used one of the candidate molecules identified in our previous study to provide a proof of principle for our approach. We combined human T21 amniocytes derived from fetuses with DS (*in vitro* model) and the Ts1Cje mouse model (*in vivo* model) to analyze the safety and efficacy of apigenin treatment. Treatment was initiated during the prenatal period when neurogenesis and synaptogenesis defects begin to occur in DS. In the Ts1Cje mouse model, we implemented a longitudinal approach to analyze the effects of prenatal apigenin treatment during different stages of the lifespan. This approach enabled us to demonstrate the safety and long-term effects of apigenin treatment. We also investigated the effects of apigenin treatment in both male and female adult Ts1Cje mice, and highlighted sex-specific specific differences in adult behavioral tests.

In this study, we only analyzed the effects of apigenin on oxidative stress and anti-oxidant capacity in T21 amniocytes, one of the many abnormalities present in individuals with DS. We used the whole transcriptome approach and showed only a mild effect of apigenin on other dysregulated pathways in T21 amniocytes. Amniocytes are a heterogeneous population of fetal stem cells, including neural stem cells that might have different responses to apigenin. We also used only one mouse model (Ts1Cje), which has a relatively mild adult phenotype, to investigate treatment effects. Our recent study (26) demonstrated fundamental differences between the three most commonly used mouse models of DS (Dp(16)1/Yey, Ts65Dn and Ts1Cje). To date, there is no consensus on what is the best mouse model that mimics closely the human condition, suggesting the use of more than one model to test the efficacy of therapeutic interventions prior to moving to a human clinical trial. Finally, we investigated the effects of treatment begun prenatally with only one (high) dose of apigenin. Using a low or medium dose of apigenin might prevent its negative effects observed in the Ts1Cje mouse model.

## FUTURE STUDIES

Given the data presented here, our future *in vitro* studies will focus on evaluating the effects of apigenin on neurogenesis and neuronal differentiation in stem cells derived from T21 and age-matched euploid cell lines. We will also investigate the mechanisms of action by which apigenin activates the PGE2/PTGER2 and HGF/MET signaling pathways and modulates G-protein signaling, cell cycle checkpoints and the immune response in DS.

Our future *in vivo* studies will also involve using additional mouse models to confirm our findings with the Ts1Cje mouse using low and medium doses of apigenin. We will also analyze the effects of apigenin on embryonic brain neurogenesis using 5-ethynyl-2′-deoxyuridine (EdU) staining. Finally, we will use the rodent version of the human CANTAB (Cambridge Neuropsychological Test Automated Battery) to evaluate highly complex cognitive functions in apigenin-treated versus untreated trisomic mice.

## CONCLUSIONS

Combining an integrated human/murine approach and the Connectivity Map database, we identified apigenin as candidate prenatal treatment for Down syndrome. We demonstrated that apigenin treatment could rescue oxidative stress/antioxidant capacity imbalance in human amniocytes from fetuses with trisomy 21, and improved several postnatal behavioral deficits in the Ts1Cje mouse model. We also demonstrated, using a whole transcriptome approach, that apigenin achieves its therapeutic action by targeting similar signaling pathways (G2/M cell cycle transition through Polo-Like kinase pathway, pro-inflammatory pathways and G-protein signaling) and up-stream regulators (PTGER2 and MET receptors, Forkhead and E2F transcription factors and IFNG, TNF, TP53 and IKBKB cell stress and inflammatory response regulators) *in vitro* and *in vivo*. These studies provide proof-of-principle that apigenin has therapeutic effects in preclinical models of Down syndrome.

## MATERIALS AND METHODS

Detailed description of the methods can be found in the “Supplementary Material”.

### In Vitro Studies on Human Amniocytes

This study was approved by the Institutional Review Boards (IRBs) at Tufts Medical Center (Protocol 5582) and Women and Infants’ Hospital (Protocol 01-0028). The amniocytes were obtained after clinically indicated prenatal karyotyping. As this was discarded material that was de-identified, patient consent was deemed unnecessary by the IRBs. Only fetal karyotype and sex were known. Second trimester amniocytes were prepared as described previously (19). The initial sample set consisted of 14 flasks of amniocytes with the following metaphase karyotypes: 47, XX, +21 (*N*=3); 47, XY, +21 (*N*=4); 46, XX (*N*=3); 46, XY (*N*=4). Gestational ages ranged from 15 3/7 to 20 2/7 weeks. Samples were matched for sex and gestational age (seven pairs were analyzed) (Supplementary Table 1 in Supplementary Material).

#### Optimal Dose Selection of Apigenin Using Cell Proliferation Assays

To determine the range of non-toxic doses that would be used to evaluate treatment efficacy, cells were either left untreated or treated with five different concentrations of apigenin (1, 2, 3, 4 and 5 µM) for three consecutive days. Automatic cell counting with using the Scepter(tm) 2.0 Cell Counter (EMD Millipore, Billerica, MA) and CellTiter 96^®^ cell proliferation assay (Promega, Madison, WI) was used. Cell proliferation in the untreated cells was normalized to 100 % and used to estimate the percent of cell proliferation in apigenin-treated cells. Toxicity was defined as doses that induced significant (*p<0.05*) or more than 15 % (even if not statistically significant) reduction of cell proliferation in both assays.

#### Oxidative Stress and Antioxidant Capacity

The level of oxidative stress damage and the effects of apigenin treatment in amniocytes from fetuses with T21 (*N*=5) and euploid fetuses (*N*=5) was quantified the Comet Assay^®^ kit according to the manufacturer’s instructions (Trevigen, Gaithersburg, MD). The percent of DNA in the “tail” versus “head” (nucleus) of the migrating cell, resembling a comet, was determined in 300-500 cells per cell line, and compared in untreated and apigenin-treated cells.

The physiological response to oxidative stress before and after apigenin treatment was measured using the OxiSelect(tm) Total Antioxidant Capacity (TAC) kit. 5,000 µg of total protein was used to evaluate TAC measured as absorbance at 490 nm and compared to a uric acid standard curve according to the manufacturer’s instructions (CellBiolabs, San Diego, CA)

#### RNA Extraction and Microarray Hybridization

Gene expression analysis was performed on treated and untreated cells using the GeneChip^®^ Human Transcriptome HT 2.0 array (Affymetrix, Santa Clara, CA). Only one apigenin dose (2 µM) was evaluated as described in the “Supplementary Material” section. Twenty-eight arrays were used (7 T21 and 7 euploid for both control and treated). Data were analyzed using a repeated-measures ANOVA that included genotype, treatment, and sample pairings between T21 and control samples. Probe sets for which either genotype or treatment was significant (at *p<0.001* and a Benjamini-Hochberg False Discovery Rate (BH-FDR) of 10%) were considered as differentially expressed genes (DEX) (99).

Pathway analyses were carried out using Database for Annotation, Visualization and Integrated Discovery (DAVID) and Ingenuity Pathway Analysis (IPA) on the top 1% up-/down-regulated genes (hereafter referred to as Marginally Expressed (MEX) Genes), and Gene Set Enrichment Analysis (GSEA) on the whole transcriptome as previously described (20). “Ingenuity Pathway Upstream Regulator Analysis” was used to identify potential upstream regulators that may be responsible for the gene expression changes observed in T21 amniocytes, and how these genes are affected after apigenin treatment. This allows the prediction of mechanism of action of apigenin and if identified upstream regulators are inhibited or activated based on a z-score algorithm (100).

### In Vivo Studies on the Ts1Cje Mouse Model

The effects of apigenin treatment were analyzed at the three different stages of murine life, embryonic, neonatal and adult (20, 26).

#### Breeding, Prenatal Apigenin Treatment and Genotyping

All murine experiments were approved by the Institutional Animal Care and Use Committee (IACUC) of Tufts University (Protocol B2013-20). Ts1Cje males (B6 T(12;16)1Cje/CjeDnJ) were crossed with C57Bl/6J females (Jackson Laboratories, Bar Harbor, ME). Breeding pairs received either purified powdered (Pow) chow F3197 (Bioserv, Flemington, NJ) or 333-400 mg/kg/day of apigenin (Apig) (2 g of apigenin in 1kg of purified powdered chow) (SelleckChem, Houston, TX). These doses were obtained using the SD=(DD x FI)/BW formula, in which SD is the single daily dose to be delivered (mg/kg/day), DD is the drug dose in the diet (2,000 mg/kg), FI is the daily food intake (5 g per mouse) and BW is the average animal weight (25-30 g) (101). Treatment was given as powdered chow with apigenin starting at the time of mating and continuing throughout pregnancy, lactation until the completion of all behavioral and biochemical studies at adulthood. Mice that received apigenin are referred to throughout the study with the subscript _Apig_, and untreated mice are referred to with the subscript _Pow_, indicating that they received only powdered chow. There was no difference in the appearance of the powdered chow versus powdered chow plus apigenin, so investigators did not know which mice had received the treatment. Genotyping and sex determination were performed by PCR using primers specific for the Ts1Cje mouse (102).

#### Embryonic Forebrain Gene Expression

For gene expression studies, total RNA was isolated from the developing forebrain using the RNA II kits following the manufacturer’s instructions (Macherey-Nagel, Bethlehem, PA). RNA was processed and hybridized on the GeneChip^®^ Mouse Gene ST 1.0 array as described previously (20). Twenty arrays were used (5 WT_Pow_, 5 Ts1Cje_Pow_, 5 WT_Apig_, 5 Ts1Cje_Apig_; 3 males and 2 females/group); each array corresponded to labeled cDNA from one sample. Analyses were performed using unpaired t-test genes. A gene with a p-value < 0.001 and a Benjamini-Hochberg False Discovery Rate (BH-FDR) of 20% was considered to be differentially-expressed (DEX).

#### Neonatal Behavior

All neonatal behavioral tests were performed blindly. The Fox scale is a general screening test used to evaluate body righting and coordination, strength, sensory maturation and extinction of rotatory behavior. Tests were performed between postnatal (P) days P3 and P21 (weaning) (103). The homing test was used to investigate olfactory-dependent spatial memory at postnatal day 12, as previously described (29). During the testing period, pups (WT_Pow_=31, Ts1Cje_Pow_=19, WT_Apig_=25, Ts1Cje_Apig_=26) were separated from the dam and placed with nesting material in a small bowl positioned on a heating pad at 37°C. A heat lamp was also placed over the mice to provide heat from above. The amount of time (latency) and presence/absence of a reflex were recorded and analyzed by a single experimenter.

#### Adult Behavior

All adult behavioral testing paradigms were performed as previously described in Aziz *et al*. (26). Exploratory behavior and locomotor activity were assessed using the open field test. Exploratory behavior was tracked during a 60-min unique trial using the Ethovision 10.5 animal tracking system (Noldus, Leesburg, VA). The total distance traveled (cm) in the center versus periphery, as well as the average velocity (cm/s), were analyzed for the treated and untreated groups. Motor coordination was investigated using the rotarod test (Med Associates, Fairfax, VT) using two different protocols (fixed speed on day 1 and accelerating speed on day 2). The time to fall was recorded in seconds and analyzed for each mouse. Hippocampal-dependent memory was analyzed using the fear conditioning test. On day one (training session), two mild foot shocks (0.5 mA for 2 s) were administered at 180 s and 240 s. On day two (testing session), mice were placed in the same chambers and the extent (or percent) of freezing, used as a measure of the animal’s memory, was analyzed as time bins of 60 s using the Freeze View software (Med Associates, Fairfax, VT).

#### Statistical Analysis

Statistical analysis was performed using the parametric t-test or two-way repeated measure ANOVA and Tukey’s multiple comparison test for normal distributions. Non-parametric Mann-Whitney and Kruskal-Wallis tests were used if values did not follow a normal distribution. For proportions (percentages) comparison of the effects of apigenin treatment on the natural history and the homing test, Chi-Square and Fisher Exact tests were used. Statistical significance was reached with a p-value < 0.05. All statistical analyses were performed using GraphPad Prism 7.03 software package. Data are presented as mean±S.D.

## Supporting information

## FIGURE LEGENDS

**Supplementary Figure 1: Effects of apigenin on cell proliferation in T21 and euploid amniocytes.** Cell proliferation was measured using two different assays and normalized to 100 % in untreated cells. Effect of apigenin were analyzed on all the cell lines together **(A)** or separated by genotype **(B)**. High doses of apigenin induced significantly reduced cell proliferation in euploid (4-5 µM) and T21 (5 µM) amniocytes. * (p<0.05), ** (p<0.01), *** (p<0.001).

**Supplementary Figure 2: Effects of apigenin on natural history and growth in Ts1Cje and WT littermates. (A-B)** Embryonic and postnatal genotype distribution in untreated and apigenin-treated mice. Genotype distribution in the untreated group followed mendelian inheritance while only 32.8% of the neonates were Ts1Cje postnatally. In the apigenin-treated group, 38% of embryos were Ts1Cje and 62% were WT. Postnatally, the genotype distribution was similar in the untreated and apigenin-treated. **(C-F)** Embryonic and postnatal growth in untreated and apigenin-treated mice. Apigenin treatment did not result in significant changes in the growth profiles in Ts1Cje and WT mice.

**Supplementary Figure 3: Effects of apigenin on early developmental milestones in Ts1Cje and WT littermates.** Untreated Ts1Cje neonates exhibit significant delays in early milestones, including surface righting **(A-B)**, cliff aversion **(C-D)**, negative geotaxis **(E-F)** and forelimb grasp **(G-H)**. Apigenin treatment partially improved Ts1Cje performance in surface righting and cliff aversion but did not affect performance in negative geotaxis and forelimb grasp tests.

**Supplementary Figure 4: Effects of apigenin on late developmental milestones in Ts1Cje and WT littermates.** Ts1Cje neonates exhibit significant delays in late milestones, including air righting **(C)**, eye opening **(D)**, ear twitch **(E)**. Apigenin treatment partially improved air righting and eye opening, but negatively affected motor development (open field).

**Supplementary Figure 5: Adverse effects of apigenin on motor coordination in untreated and apigenin-treated adult Ts1Cje and WT males and females.** Motor coordination was analyzed using the rotarod test in untreated and apigenin-treated WT and Ts1Cje males (WT_Pow_=13, Ts1Cje_Pow_=12, WT_Apig_=17, Ts1Cje_Apig_=16) and females (WT_Pow_=8, Ts1Cje_Pow_=7, WT_Apig_=14, Ts1Cje_Apig_=12). **(A-B)** Performance of untreated and apigenin-treated Ts1Cje and WT male and female mice in the static speed trial at 32 RPM. **(C-D)** Performance of untreated and apigenin-treated Ts1Cje and WT male and female mice in the accelerating speed trial (4-40 RPM). Ts1Cje males exhibited significant motor coordination deficits (fell off the rotarod faster) compared to their WT littermates, however, motor coordination was not significantly affected in Ts1Cje female mice. Apigenin treatment had negative effects on motor coordination in both WT and Ts1Cje male and female mice. * (p<0.05), ** (p<0.01), *** (p<0.001).

## SUPPLEMENTARY TABLES

**Supplementary Table 1: Karyotype and gestational age information of the human trisomy 21 (T21) and euploid amniocytes pairs used in this study.**

**Supplementary Table 2: Effects of apigenin on differentially expressed (DEX) genes in age and sex-matched T21 and euploid amniocytes. (2A)** List of DEX genes in T21 versus euploid amniocytes. **(2B)** Regulatory effects of apigenin treatment on chromosome 21 genes in T21 amniocytes. **(2C)** List of DEX genes induced by apigenin treatment in T21 amniocytes.

**Supplementary Table 3: Marginally expressed (MEX) genes (Top 1% up-and down-regulated genes) in untreated and apigenin-treated T21 and euploid amniocytes. (3A)** List of MEX genes specifically up- and down-regulated after apigenin treatment. **(3B)** List of MEX genes in untreated human T21 versus euploid amniocytes. **(3C)** List of MEX genes in apigenin-treated T21 compared to untreated T21 amniocytes. **(3D)** List of MEX genes in apigenin-treated T21 amniocytes compared to untreated euploid amniocytes. **(3E)** List of MEX genes in apigenin-treated euploid compared to untreated euploid amniocytes. Up-regulated genes are highlighted in “Red” and down-regulated genes in “Blue” and the percent of gene expression change is indicated for each comparison.

**Supplementary Table 4: Summary of DAVID dysregulated pathways in untreated and apigenin-treated T21 and euploid amniocytes. (4A)** DAVID dysregulated pathways in untreated human T21 versus euploid amniocytes. **(4B)** DAVID dysregulated pathways in apigenin-treated T21 compared to untreated T21 amniocytes. **(4C)** DAVID dysregulated pathways in apigenin-treated T21 amniocytes compared to untreated euploid amniocytes. **(4D)** DAVID dysregulated pathways in apigenin-treated euploid compared to untreated euploid amniocytes.

**Supplementary Table 5: Summary of GSEA dysregulated pathways in untreated and apigenin-treated T21 and euploid amniocytes. (5A)** GSEA dysregulated pathways in untreated human T21 versus euploid amniocytes. **(5B)** GSEA dysregulated pathways in apigenin-treated T21 compared to untreated T21 amniocytes. **(5C)** GSEA dysregulated pathways in apigenin-treated T21 amniocytes compared to untreated euploid amniocytes. **(5D)** GSEA dysregulated pathways in apigenin-treated euploid compared to untreated euploid amniocytes.

**Supplementary Table 6: Summary of IPA dysregulated pathways in untreated and apigenin-treated T21 and euploid amniocytes. (6A)** IPA dysregulated pathways in untreated human T21 versus euploid amniocytes. **(6B)** IPA dysregulated pathways in apigenin-treated T21 compared to untreated T21 amniocytes. **(6C)** IPA dysregulated pathways in apigenin-treated T21 amniocytes compared to untreated euploid amniocytes. **(6D)** IPA dysregulated pathways in apigenin-treated euploid compared to untreated euploid amniocytes.

**Supplementary Table 7: Summary of IPA predicted upstream regulators in untreated and apigenin-treated T21 amniocytes. (7A)** IPA predicted upstream regulators in untreated human T21 versus euploid amniocytes. **(7B)** in apigenin-treated T21 compared to untreated T21 amniocytes. **(7C)** IPA predicted upstream regulators in apigenin-treated T21 amniocytes compared to untreated euploid amniocytes. **(7D)** IPA predicted upstream regulators in apigenin-treated euploid compared to untreated euploid amniocytes.

**Supplementary Table 8: Effects of apigenin on DEX genes in Ts1Cje and WT E15.5 forebrain. (8A)** Regulatory effects of apigenin treatment on the DEX genes in untreated Ts1Cje E15.5 forebrain. **(8B)** DEX Genes in apigenin-treated Ts1Cje compared to apigenin-treated E15.5 forebrain. **(8C)** DEX genes in apigenin-treated Ts1Cje and WT E15.5 forebrain compared to their untreated counterparts.

**Supplementary Table 9: MEX genes in untreated and apigenin-treated Ts1Cje and WT E15.5 forebrain. (9A)** MEX genes in untreated Ts1Cje compared to untreated WT embryonic forebrain. **(9B)** MEX genes in apigenin-treated WT compared to untreated WT embryonic forebrain. **(9C)** MEX genes in apigenin-treated Ts1Cje compared to untreated WT embryonic forebrain. **(9D)** MEX genes in apigenin-treated Ts1Cje compared to apigenin-treated WT embryonic forebrain. **(9E)** MEX genes in apigenin-treated Ts1Cje compared to untreated Ts1Cje embryonic forebrain. Up-regulated genes are highlighted in “Red” and down-regulated genes in “Blue” and the percent of expression change is indicated for each comparison.

**Supplementary Table 10: Summary of DAVID dysregulated pathways in untreated and apigenin-treated Ts1Cje and WT E15.5 forebrain. (10A)** DAVID dysregulated pathways in Ts1Cje compared to untreated WT embryonic forebrain. **(10B)** DAVID dysregulated pathways in apigenin-treated Ts1Cje compared to untreated Ts1Cje embryonic forebrain. **(10C)** DAVID dysregulated pathways in apigenin-treated Ts1Cje compared to untreated WT embryonic forebrain. **(10D)** DAVID dysregulated pathways in apigenin-treated WT compared to untreated WT embryonic forebrain.

**Supplementary Table 11: Summary of GSEA dysregulated pathways in untreated and apigenin-treated Ts1Cje and WT E15.5 forebrain. (11A)** GSEA dysregulated pathways in Ts1Cje compared to untreated WT embryonic forebrain. **(11B)** GSEA dysregulated pathways in apigenin-treated Ts1Cje compared to untreated Ts1Cje embryonic forebrain. **(11C)** GSEA dysregulated pathways in apigenin-treated Ts1Cje compared to untreated WT embryonic forebrain. **(11D)** GSEA dysregulated pathways in apigenin-treated WT compared to untreated WT embryonic forebrain.

**Supplementary Table 12: Summary of IPA dysregulated pathways in untreated and apigenin-treated Ts1Cje and WT E15.5 forebrain. (12A)** IPA dysregulated pathways in Ts1Cje compared to untreated WT embryonic forebrain. **(12B)** IPA dysregulated pathways in apigenin-treated Ts1Cje compared to untreated Ts1Cje embryonic forebrain. **(12C)** IPA dysregulated pathways in apigenin-treated Ts1Cje compared to untreated WT embryonic forebrain. **(12D)** IPA dysregulated pathways in apigenin-treated WT compared to untreated WT embryonic forebrain.

**Supplementary Table 13: Summary of IPA predicted upstream regulators in untreated and apigenin-treated Ts1Cje and WT E15.5 forebrain. (13A)** IPA predicted upstream regulators in Ts1Cje compared to untreated WT embryonic forebrain. **(13B)** IPA predicted upstream regulators in apigenin-treated Ts1Cje compared to untreated Ts1Cje embryonic forebrain. **(13C)** IPA predicted upstream regulators in apigenin-treated Ts1Cje compared to untreated WT embryonic forebrain. **(13D)** IPA predicted upstream regulators in apigenin-treated WT compared to untreated WT embryonic forebrain.

**Supplementary Table 14: Comparison of the main upstream regulators predicted to be activated or inhibited by apigenin in human T21 amniocytes and Ts1Cje mouse model.**

## AUTHOR CONTRIBUTIONS

**Conceptualization:** F.G., D.W.B.

**Methodology:** F.G., J.L.A.P., A.E.S, L.J.M, F.A., D.W.B.

**Validation:** F.G., J.L.A.P., D.W.B.

**Formal analysis:** F.G., J.L.A.P., A.E.S, F.A

**Investigation:** F.G., J.L.A.P., A.E.S, F.A

**Resources:** D.W.B.

**Writing:** – original draft F.G., D.W.B.

**Writing - review & editing:** F.G., J.L.A.P., A.E.S, F.A, D.W.B.

**Supervision:** F.G., D.W.B

**Funding acquisition:** F.G., D.W.B.

## ACKNOWLEDGMENTS

This study was funded by the National Institutes of Health (NICHD R01HD058880) and a sponsored research grant from Verinata Health, Inc., an Illumina company.

The authors also wish to acknowledge the help and support of Donna Slonim Ph.D and the Tufts University Neuroscience Behavior Core Facility.

## DATA AVAILABILITY

All gene expression data have been deposited into the Gene Expression Omnibus (GEO). GSE identifiers will be added when received from GEO.

## REFERENCES

1. S. Taylor-Phillips, K. Freeman, J. Geppert, A. Agbebiyi, O. A. Uthman, J. Madan, A. Clarke, S. Quenby, A. Clarke, Accuracy of non-invasive prenatal testing using cell-free DNA for detection of Down, Edwards and Patau syndromes: a systematic review and meta-analysis. BMJ Open 6, e010002 (2016).

2. S. J. Ralston, D. Wertz, D. Chelmow, S. D. Craigo, D. W. Bianchi, Pregnancy outcomes after prenatal diagnosis of aneuploidy. Obstet Gynecol 97, 729–733 (2001).

3. F. Guedj, D. W. Bianchi, Noninvasive prenatal testing creates an opportunity for antenatal treatment of Down syndrome. Prenat Diagn 33, 614–618 (2013).

4. D. W. Bianchi, From prenatal genomic diagnosis to fetal personalized medicine: progress and challenges. Nat Med 18, 1041–1051 (2012).

5. B. Schmidt-Sidor, K. E. Wisniewski, T. H. Shepard, E. A. Sersen, Brain growth in Down syndrome subjects 15 to 22 weeks of gestational age and birth to 60 months. Clin Neuropathol 9, 181–190 (1990).

6. S. Guidi, P. Bonasoni, C. Ceccarelli, D. Santini, F. Gualtieri, E. Ciani, R. Bartesaghi, Neurogenesis impairment and increased cell death reduce total neuron number in the hippocampal region of fetuses with Down syndrome. Brain Pathol 18, 180–197 (2008).

7. S. Guidi, E. Ciani, P. Bonasoni, D. Santini, R. Bartesaghi, Widespread proliferation impairment and hypocellularity in the cerebellum of fetuses with Down syndrome. Brain Pathol 21, 361–373 (2011).

8. K. E. Wisniewski, B. Schmidt-Sidor, Postnatal delay of myelin formation in brains from Down syndrome infants and children. Clin Neuropathol 8, 55–62 (1989).

9. H. Abraham, A. Vincze, B. Veszpremi, A. Kravjak, E. Gomori, G. G. Kovacs, L. Seress, Impaired myelination of the human hippocampal formation in Down syndrome. Int J Dev Neurosci 30, 147–158 (2012).

10. P. G. Hepper, S. Shahidullah, Habituation in normal and Down’s syndrome fetuses. Q J Exp Psychol B 44, 305–317 (1992).

11. F. Guedj, D. W. Bianchi, J. M. Delabar, Prenatal treatment of Down syndrome: a reality? Curr Opin Obstet Gynecol 26, 92–103 (2014).

12. F. Stagni, A. Giacomini, S. Guidi, E. Ciani, R. Bartesaghi, Timing of therapies for Down syndrome: the sooner, the better. Front Behav Neurosci 9, 265 (2015).

13. K. J. Gardiner, Pharmacological approaches to improving cognitive function in Down syndrome: current status and considerations. Drug Des Devel Ther 9, 103–125 (2014).

14. D. Rice, S. Barone, Jr., Critical periods of vulnerability for the developing nervous system: evidence from humans and animal models. Environ Health Perspect 108 Suppl 3, 511–533 (2000).

15. R. M. Meredith, Sensitive and critical periods during neurotypical and aberrant neurodevelopment: a framework for neurodevelopmental disorders. Neurosci Biobehav Rev 50, 180–188 (2015).

16. S. Guidi, F. Stagni, P. Bianchi, E. Ciani, A. Giacomini, M. De Franceschi, R. Moldrich, N. Kurniawan, K. Mardon, A. Giuliani, L. Calza, R. Bartesaghi, Prenatal pharmacotherapy rescues brain development in a Down’s syndrome mouse model. Brain 137, 380–401 (2014).

17. M. Incerti, K. Horowitz, R. Roberson, D. Abebe, L. Toso, M. Caballero, C. Y. Spong, Prenatal treatment prevents learning deficit in Down syndrome model. PLoS One 7, e50724 (2012).

18. J. A. Ash, R. Velazquez, C. M. Kelley, B. E. Powers, S. D. Ginsberg, E. J. Mufson, B. J. Strupp, Maternal choline supplementation improves spatial mapping and increases basal forebrain cholinergic neuron number and size in aged Ts65Dn mice. Neurobiol Dis 70, 32–42 (2014).

19. D. K. Slonim, K. Koide, K. L. Johnson, U. Tantravahi, J. M. Cowan, Z. Jarrah, D. W. Bianchi, Functional genomic analysis of amniotic fluid cell-free mRNA suggests that oxidative stress is significant in Down syndrome fetuses. Proc Natl Acad Sci U S A 106, 9425–9429 (2009).

20. F. Guedj, J. L. Pennings, L. J. Massingham, H. C. Wick, A. E. Siegel, U. Tantravahi, D. W. Bianchi, An integrated human/murine transcriptome and pathway approach to identify prenatal treatments for Down syndrome. Sci Rep 6, 32353 (2016).

21. A. J. Lamb, E. D. Crawford, D. Peck, J. W. Modell, I. C. Blat, M. J. Wrobel, J. Lerner, J. P. Brunet, A. Subramanian, K. N. Ross, M. Reich, H. Hieronymus, G. Wei, S. A. Armstrong, S. J. Haggarty, P. A. Clemons, R. Wei, S. A. Carr, E. S. Lander, T. R. Golub, The Connectivity Map: A. using gene-expression signatures to connect small molecules, genes, and disease. Science 313, 1929–1935 (2006).

22. M. Venigalla, E. Gyengesi, G. Munch, Curcumin and Apigenin - novel and promising therapeutics against chronic neuroinflammation in Alzheimer’s disease. Neural Regen Res 10, 1181–1185 (2015).

23. M. Venigalla, S. Sonego, E. Gyengesi, M. J. Sharman, G. Munch, Novel promising therapeutics against chronic neuroinflammation and neurodegeneration in Alzheimer’s disease. Neurochem Int 95, 63–74 (2016).

24. J. Kalivarathan, S. P. Chandrasekaran, K. Kalaivanan, V. Ramachandran, A. Carani Venkatraman, Apigenin attenuates hippocampal oxidative events, inflammation and pathological alterations in rats fed high fat, fructose diet. Biomed Pharmacother 89, 323–331 (2017).

25. K. Rezai-Zadeh, J. Ehrhart, Y. Bai, P. R. Sanberg, P. Bickford, J. Tan, R. D. Shytle, Apigenin and luteolin modulate microglial activation via inhibition of STAT1-induced CD40 expression. J Neuroinflammation 5, 41 (2008).

26. N. M. Aziz, F. Guedj, J. L. A. Pennings, J. L. Olmos-Serrano, A. Siegel, T. F. Haydar, D. W. Bianchi, Lifespan analysis of brain development, gene expression and behavioral phenotypes in the Ts1Cje, Ts65Dn and Dp(16)1/Yey mouse models of Down syndrome. Dis Model Mech, Jun 12; 11(6).pii.dmm031013.doi:10.1242/dmm.031013 (2018).

27. K. D. Sullivan, H. C. Lewis, A. A. Hill, A. Pandey, L. P. Jackson, J. M. Cabral, K. P. Smith, L. A. Liggett, E. B. Gomez, M. D. Galbraith, J. DeGregori, J. M. Espinosa, Trisomy 21 consistently activates the interferon response. Elife, Jul 29;5.pii.e16220.doi:10.7554/eLife.16220 (2016).

28. L. Zhao, J. L. Wang, R. Liu, X. X. Li, J. F. Li, L. Zhang, Neuroprotective, anti-amyloidogenic and neurotrophic effects of apigenin in an Alzheimer’s disease mouse model. Molecules 18, 9949–9965 (2013).

29. F. Guedj, J. L. Pennings, M. A. Ferres, L. C. Graham, H. C. Wick, K. A. Miczek, D. K. Slonim, D. W. Bianchi, The fetal brain transcriptome and neonatal behavioral phenotype in the Ts1Cje mouse model of Down syndrome. Am J Med Genet A 167A, 1993–2008 (2015).

30. E. Barone, A. Arena, E. Head, D. A. Butterfield, M. Perluigi, Disturbance of redox homeostasis in Down Syndrome: Role of iron dysmetabolism. Free Radic Biol Med 114, 84–93 (2018).

31. J. Muchova, I. Zitnanova, Z. Durackova, Oxidative stress and Down syndrome. Do antioxidants play a role in therapy? Physiol Res 63, 535–542 (2014).

32. G. Pagano, G. Castello, Oxidative stress and mitochondrial dysfunction in Down syndrome. Adv Exp Med Biol 724, 291–299 (2012).

33. I. T. Lott, Antioxidants in Down syndrome. Biochim Biophys Acta 1822, 657–663 (2012).

34. R. C. Iannello, P. J. Crack, J. B. de Haan, I. Kola, Oxidative stress and neural dysfunction in Down syndrome. J Neural Transm Suppl 57, 257–267 (1999).

35. A. Gimeno, J. L. Garcia-Gimenez, L. Audi, N. Toran, P. Andaluz, F. Dasi, J. Vina, F. V. Pallardo, Decreased cell proliferation and higher oxidative stress in fibroblasts from Down Syndrome fetuses. Preliminary study. Biochim Biophys Acta 1842, 116–125 (2014).

36. M. Perluigi, F. di Domenico, A. Fiorini, A. Cocciolo, A. Giorgi, C. Foppoli, D. A. Butterfield, M. Giorlandino, C. Giorlandino, M. E. Schinina, R. Coccia, Oxidative stress occurs early in Down syndrome pregnancy: A redox proteomics analysis of amniotic fluid. Proteomics Clin Appl 5, 167–178 (2011).

37. J. B. de Haan, B. Susil, M. Pritchard, I. Kola, An altered antioxidant balance occurs in Down syndrome fetal organs: implications for the “gene dosage effect” hypothesis. J Neural Transm Suppl, 67–83 (2003).

38. H. Sharma, R. Kanwal, N. Bhaskaran, S. Gupta, Plant flavone apigenin binds to nucleic acid bases and reduces oxidative DNA damage in prostate epithelial cells. PLoS One 9, e91588 (2014).

39. S. S. Kang, J. Y. Lee, Y. K. Choi, G. S. Kim, B. H. Han, Neuroprotective effects of flavones on hydrogen peroxide-induced apoptosis in SH-SY5Y neuroblostoma cells. Bioorg Med Chem Lett 14, 2261–2264 (2004).

40. F. An, X. Cao, H. Qu, S. Wang, Attenuation of oxidative stress of erythrocytes by the plant-derived flavonoids vitexin and apigenin. Pharmazie 70, 724–732 (2015).

41. S. F. Nabavi, H. Khan, G. D’Onofrio, D. Samec, S. Shirooie, A. R. Dehpour, S. Arguelles, S. Habtemariam, E. Sobarzo-Sanchez, Apigenin as neuroprotective agent: Of mice and men. Pharmacol Res 128, 359–365 (2018).

42. B. J. Jeon, H. M. Yang, Y. S. Lyu, H. O. Pae, S. M. Ju, B. H. Jeon, Apigenin inhibits indoxyl sulfate-induced endoplasmic reticulum stress and anti-proliferative pathways, CHOP and IL-6/p21, in human renal proximal tubular cells. Eur Rev Med Pharmacol Sci 19, 2303–2310 (2015).

43. P. S. Wu, J. H. Yen, M. C. Kou, M. J. Wu, Luteolin and Apigenin Attenuate 4-Hydroxy-2-Nonenal-Mediated Cell Death through Modulation of UPR, Nrf2-ARE and MAPK Pathways in PC12 Cells. PLoS One 10, e0130599 (2015).

44. Y. M. Lee, G. Lee, T. I. Oh, B. M. Kim, D. W. Shim, K. H. Lee, Y. J. Kim, B. O. Lim, J. H. Lim, Inhibition of glutamine utilization sensitizes lung cancer cells to apigenin-induced apoptosis resulting from metabolic and oxidative stress. Int J Oncol 48, 399–408 (2016).

45. K. Banerjee, M. Mandal, Oxidative stress triggered by naturally occurring flavone apigenin results in senescence and chemotherapeutic effect in human colorectal cancer cells. Redox Biol 5, 153–162 (2015).

46. V. Stepanic, A. C. Gasparovic, K. G. Troselj, D. Amic, N. Zarkovic, Selected attributes of polyphenols in targeting oxidative stress in cancer. Curr Top Med Chem 15, 496–509 (2015).

47. R. P. Souza, P. S. Bonfim-Mendonca, F. Gimenes, B. A. Ratti, V. Kaplum, M. L. Bruschi, C. V. Nakamura, S. O. Silva, S. S. Maria-Engler, M. E. Consolaro, Oxidative Stress Triggered by Apigenin Induces Apoptosis in a Comprehensive Panel of Human Cervical Cancer-Derived Cell Lines. Oxid Med Cell Longev 2017, 1512745 (2017).

48. J. M. M. Salmani, X. P. Zhang, J. A. Jacob, B. A. Chen, Apigenin’s anticancer properties and molecular mechanisms of action: Recent advances and future prospectives. Chin J Nat Med 15, 321–329 (2017).

49. D. L. McKay, J. B. Blumberg, A review of the bioactivity and potential health benefits of chamomile tea (Matricaria recutita L.). Phytother Res 20, 519–530 (2006).

50. F. Qi, Q. Chen, H. Chen, H. Yan, B. Chen, X. Xiang, C. Liang, Q. Yi, M. Zhang, H. Cheng, Z. Zhang, J. Huang, F. Wang, WAC Promotes Polo-like Kinase 1 Activation for Timely Mitotic Entry. Cell Rep 24, 546–556 (2018).

51. L. Gheghiani, D. Loew, B. Lombard, J. Mansfeld, O. Gavet, PLK1 Activation in Late G2 Sets Up Commitment to Mitosis. Cell Rep 19, 2060–2073 (2017).

52. T. H. Tseng, M. H. Chien, W. L. Lin, Y. C. Wen, J. M. Chow, C. K. Chen, T. C. Kuo, W. J. Lee, Inhibition of MDA-MB-231 breast cancer cell proliferation and tumor growth by apigenin through induction of G2/M arrest and histone H3 acetylation-mediated p21(WAF1/CIP1) expression. Environ Toxicol 32, 434–444 (2017).

53. J. Cai, X. L. Zhao, A. W. Liu, H. Nian, S. H. Zhang, Apigenin inhibits hepatoma cell growth through alteration of gene expression patterns. Phytomedicine 18, 366–373 (2011).

54. Q. Zhang, X. H. Zhao, Z. J. Wang, Cytotoxicity of flavones and flavonols to a human esophageal squamous cell carcinoma cell line (KYSE-510) by induction of G2/M arrest and apoptosis. Toxicol In Vitro 23, 797–807 (2009).

55. M. B. Ujiki, X. Z. Ding, M. R. Salabat, D. J. Bentrem, L. Golkar, B. Milam, M. S. Talamonti, R. H. Bell, Jr., T. Iwamura, T. E. Adrian, Apigenin inhibits pancreatic cancer cell proliferation through G2/M cell cycle arrest. Mol Cancer 5, 76 (2006).

56. M. M. Corsi, G. Dogliotti, F. Pedroni, E. Palazzi, P. Magni, M. Chiappelli, F. Licastro, Plasma nerve growth factor (NGF) and inflammatory cytokines (IL-6 and MCP-1) in young and adult subjects with Down syndrome: an interesting pathway. Neuro Endocrinol Lett 27, 773–778 (2006).

57. F. Licastro, M. Chiappelli, E. Porcellini, M. Trabucchi, A. Marocchi, M. M. Corsi, Altered vessel signalling molecules in subjects with Downs syndrome. Int J Immunopathol Pharmacol 19, 181–185 (2006).

58. M. F. Iulita, A. Ower, C. Barone, R. Pentz, P. Gubert, C. Romano, R. A. Cantarella, F. Elia, S. Buono, M. Recupero, C. Romano, S. Castellano, P. Bosco, S. Di Nuovo, F. Drago, F. Caraci, A. C. Cuello, An inflammatory and trophic disconnect biomarker profile revealed in Down syndrome plasma: Relation to cognitive decline and longitudinal evaluation. Alzheimers Dement 12, 1132–1148 (2016).

59. X. Zhang, G. Wang, E. C. Gurley, H. Zhou, Flavonoid apigenin inhibits lipopolysaccharide-induced inflammatory response through multiple mechanisms in macrophages. PLoS One 9, e107072 (2014).

60. J. Hong, A. Fristiohady, C. H. Nguyen, D. Milovanovic, N. Huttary, S. Krieger, J. Hong, S. Geleff, P. Birner, W. Jager, A. Ozmen, L. Krenn, G. Krupitza, Apigenin and Luteolin Attenuate the Breaching of MDA-MB231 Breast Cancer Spheroids Through the Lymph Endothelial Barrier in Vitro. Front Pharmacol 9, 220 (2018).

61. D. Bauer, N. Redmon, E. Mazzio, K. F. Soliman, Apigenin inhibits TNFalpha/IL-1alpha-induced CCL2 release through IKBK-epsilon signaling in MDA-MB-231 human breast cancer cells. PLoS One 12, e0175558 (2017).

62. Y. Saito, A. Oka, M. Mizuguchi, K. Motonaga, Y. Mori, L. E. Becker, K. Arima, J. Miyauchi, S. Takashima, The developmental and aging changes of Down’s syndrome cell adhesion molecule expression in normal and Down’s syndrome brains. Acta Neuropathol 100, 654–664 (2000).

63. G. M. Barlow, B. Micales, G. E. Lyons, J. R. Korenberg, Down syndrome cell adhesion molecule is conserved in mouse and highly expressed in the adult mouse brain. Cytogenet Cell Genet 94, 155–162 (2001).

64. A. Cooper, G. Grigoryan, L. Guy-David, M. M. Tsoory, A. Chen, E. Reuveny, Trisomy of the G protein-coupled K+ channel gene, Kcnj6, affects reward mechanisms, cognitive functions, and synaptic plasticity in mice. Proc Natl Acad Sci U S A 109, 2642–2647 (2012).

65. X. Jiang, C. Liu, T. Yu, L. Zhang, K. Meng, Z. Xing, P. V. Belichenko, A. M. Kleschevnikov, A. Pao, J. Peresie, S. Wie, W. C. Mobley, Y. E. Yu, Genetic dissection of the Down syndrome critical region. Hum Mol Genet 24, 6540–6551 (2015).

66. M. Raveau, T. Nakahari, S. Asada, K. Ishihara, K. Amano, A. Shimohata, H. Sago, K. Yamakawa, Brain ventriculomegaly in Down syndrome mice is caused by Pcp4 dose-dependent cilia dysfunction. Hum Mol Genet 26, 923–931 (2017).

67. F. Mouton-Liger, I. Sahun, T. Collin, P. Lopes Pereira, D. Masini, S. Thomas, E. Paly, S. Luilier, S. Meme, Q. Jouhault, S. Bennai, J. C. Beloeil, J. C. Bizot, Y. Herault, M. Dierssen, N. Creau, Developmental molecular and functional cerebellar alterations induced by PCP4/PEP19 overexpression: implications for Down syndrome. Neurobiol Dis 63, 92–106 (2014).

68. C. Anusha, T. Sumathi, L. D. Joseph, Protective role of apigenin on rotenone induced rat model of Parkinson’s disease: Suppression of neuroinflammation and oxidative stress mediated apoptosis. Chem Biol Interact 269, 67–79 (2017).

69. A. T. Smolinski, J. J. Pestka, Modulation of lipopolysaccharide-induced proinflammatory cytokine production in vitro and in vivo by the herbal constituents apigenin (chamomile), ginsenoside Rb(1) (ginseng) and parthenolide (feverfew). Food Chem Toxicol 41, 1381–1390 (2003).

70. S. P. Patil, P. D. Jain, J. S. Sancheti, P. J. Ghumatkar, R. Tambe, S. Sathaye, Neuroprotective and neurotrophic effects of Apigenin and Luteolin in MPTP induced parkinsonism in mice. Neuropharmacology 86, 192–202 (2014).

71. L. Chen, W. Xie, W. Xie, W. Zhuang, C. Jiang, N. Liu, Apigenin attenuates isoflurane-induced cognitive dysfunction via epigenetic regulation and neuroinflammation in aged rats. Arch Gerontol Geriatr 73, 29–36 (2017).

72. D. Minc-Golomb, H. Knobler, Y. Groner, Gene dosage of CuZnSOD and Down’s syndrome: diminished prostaglandin synthesis in human trisomy 21, transfected cells and transgenic mice. EMBO J 10, 2119–2124 (1991).

73. A. Oka, S. Takashima, Induction of cyclo-oxygenase 2 in brains of patients with Down’s syndrome and dementia of Alzheimer type: specific localization in affected neurons and axons. Neuroreport 8, 1161–1164 (1997).

74. M. Barr-Agholme, L. Krekmanova, T. Yucel-Lindberg, K. Shinoda, T. Modeer, Prostaglandin E2 level in gingival crevicular fluid from patients with Down syndrome. Acta Odontol Scand 55, 101–105 (1997).

75. Y. Otsuka, M. Ito, M. Yamaguchi, S. Saito, K. Uesu, K. Kasai, Y. Abiko, J. Mega, Enhancement of lipopolysaccharide-stimulated cyclooxygenase-2 mRNA expression and prostaglandin E2 production in gingival fibroblasts from individuals with Down syndrome. Mech Ageing Dev 123, 663–674 (2002).

76. J. Jiang, R. Dingledine, Prostaglandin receptor EP2 in the crosshairs of anti-inflammation, anti-cancer, and neuroprotection. Trends Pharmacol Sci 34, 413–423 (2013).

77. M. Prat, F. Oltolina, C. Basilico, Monoclonal Antibodies against the MET/HGF Receptor and Its Ligand: Multitask Tools with Applications from Basic Research to Therapy. Biomedicines 2, 359–383 (2014).

78. T. Nakamura, S. Mizuno, The discovery of hepatocyte growth factor (HGF) and its significance for cell biology, life sciences and clinical medicine. Proc Jpn Acad Ser B Phys Biol Sci 86, 588–610 (2010).

79. J. W. Wright, J. W. Harding, The Brain Hepatocyte Growth Factor/c-Met Receptor System: A New Target for the Treatment of Alzheimer’s Disease. J Alzheimers Dis 45, 985–1000 (2015).

80. T. W. Wang, H. Zhang, M. R. Gyetko, J. M. Parent, Hepatocyte growth factor acts as a mitogen and chemoattractant for postnatal subventricular zone-olfactory bulb neurogenesis. Mol Cell Neurosci 48, 38–50 (2011).

81. S. Mohan, S. Narumiya, S. Dore, Neuroprotective role of prostaglandin PGE2 EP2 receptor in hemin-mediated toxicity. Neurotoxicology 46, 53–59 (2015).

82. D. Liu, L. Wu, R. Breyer, M. P. Mattson, K. Andreasson, Neuroprotection by the PGE2 EP2 receptor in permanent focal cerebral ischemia. Ann Neurol 57, 758–761 (2005).

83. T. J. Montine, D. Milatovic, R. C. Gupta, T. Valyi-Nagy, J. D. Morrow, R. M. Breyer, Neuronal oxidative damage from activated innate immunity is EP2 receptor-dependent. J Neurochem 83, 463–470 (2002).

84. H. Yang, J. Zhang, R. M. Breyer, C. Chen, Altered hippocampal long-term synaptic plasticity in mice deficient in the PGE2 EP2 receptor. J Neurochem 108, 295–304 (2009).

85. A. Savonenko, P. Munoz, T. Melnikova, Q. Wang, X. Liang, R. M. Breyer, T. J. Montine, A. Kirkwood, K. Andreasson, Impaired cognition, sensorimotor gating, and hippocampal long-term depression in mice lacking the prostaglandin E2 EP2 receptor. Exp Neurol 217, 63–73 (2009).

86. N. S. Woodling, K. I. Andreasson, Untangling the Web: Toxic and Protective Effects of Neuroinflammation and PGE2 Signaling in Alzheimer’s Disease. ACS Chem Neurosci 7, 454–463 (2016).

87. P. Sharma, S. Sharma, D. Singh, Apigenin reverses behavioral impairments and cognitive decline in kindled mice via CREB-BDNF upregulation in the hippocampus. Nutr Neurosci, 1–10 (2018).

88. B. K. Vazhayil, S. S. Rajagopal, T. Thangavelu, G. Swaminathan, E. Rajagounder, Neuroprotective effect of Clerodendrum serratum Linn. leaves extract against acute restraint stress-induced depressive-like behavioral symptoms in adult mice. Indian J Pharmacol 49, 34–41 (2017).

89. H. Viola, C. Wasowski, M. Levi de Stein, C. Wolfman, R. Silveira, F. Dajas, J. H. Medina, A. C. Paladini, Apigenin, a component of Matricaria recutita flowers, is a central benzodiazepine receptors-ligand with anxiolytic effects. Planta Med 61, 213–216 (1995).

90. R. Crupi, I. Paterniti, A. Ahmad, M. Campolo, E. Esposito, S. Cuzzocrea, Effects of palmitoylethanolamide and luteolin in an animal model of anxiety/depression. CNS Neurol Disord Drug Targets 12, 989–1001 (2013).

91. F. S. Tsai, H. Y. Cheng, M. T. Hsieh, C. R. Wu, Y. C. Lin, W. H. Peng, The ameliorating effects of luteolin on beta-amyloid-induced impairment of water maze performance and passive avoidance in rats. Am J Chin Med 38, 279–291 (2010).

92. F. Tu, Q. Pang, T. Huang, Y. Zhao, M. Liu, X. Chen, Apigenin Ameliorates Post-Stroke Cognitive Deficits in Rats Through Histone Acetylation-Mediated Neurochemical Alterations. Med Sci Monit 23, 4004–4013 (2017).

93. X. Fu, J. Zhang, L. Guo, Y. Xu, L. Sun, S. Wang, Y. Feng, L. Gou, L. Zhang, Y. Liu, Protective role of luteolin against cognitive dysfunction induced by chronic cerebral hypoperfusion in rats. Pharmacol Biochem Behav 126, 122–130 (2014).

94. H. Wang, H. Wang, H. Cheng, Z. Che, Ameliorating effect of luteolin on memory impairment in an Alzheimer’s disease model. Mol Med Rep 13, 4215–4220 (2016).

95. T. Y. Lin, C. W. Lu, S. J. Wang, Luteolin protects the hippocampus against neuron impairments induced by kainic acid in rats. Neurotoxicology 55, 48–57 (2016).

96. K. Hara, Y. Haranishi, T. Terada, Y. Takahashi, M. Nakamura, T. Sata, Effects of intrathecal and intracerebroventricular administration of luteolin in a rat neuropathic pain model. Pharmacol Biochem Behav 125, 78–84 (2014).

97. A. Gradolatto, J. P. Basly, R. Berges, C. Teyssier, M. C. Chagnon, M. H. Siess, M. C. Canivenc-Lavier, Pharmacokinetics and metabolism of apigenin in female and male rats after a single oral administration. Drug Metab Dispos 33, 49–54 (2005).

98. T. Lorand, E. Vigh, J. Garai, Hormonal action of plant derived and anthropogenic non-steroidal estrogenic compounds: phytoestrogens and xenoestrogens. Curr Med Chem 17, 3542–3574 (2010).

99. Benjamini Y, Hochberg Y. Controlling the false discovery rate: a practical and powerful approach to multiple testing. J R Stat Soc B 57, 289–300 (1995).

100. A. Kramer, J. Green, J. Pollard, Jr., S. Tugendreich, Causal analysis approaches in Ingenuity Pathway Analysis. Bioinformatics 30, 523–530 (2014).

101. Ricci, M. Dosing animal via diet: This can be a simple and efficient method for delivering compounds to lab animals when done correctly. ALN World. 5, 1–4 (2012).

102. M. A. Ferres, D. W. Bianchi, A. E. Siegel, R. T. Bronson, G. S. Huggins, F. Guedj, Perinatal Natural History of the Ts1Cje Mouse Model of Down Syndrome: Growth Restriction, Early Mortality, Heart Defects, and Delayed Development. PLoS One 11, e0168009 (2016).

103. Hill JM, Lim MA, Stone MM. Developmental Milestones in the Newborn Mouse. Neuromethods. 39: 131–149 (2008).

104. N. E. Wilsher, R. R. Arroo, M. T. Matsoukas, A. M. Tsatsakis, D. A. Spandidos, V. P. Androutsopoulos, Cytochrome P450 CYP1 metabolism of hydroxylated flavones and flavonols: Selective bioactivation of luteolin in breast cancer cells. Food Chem Toxicol 110, 383–394 (2017).

105. V. P. Androutsopoulos, A. Papakyriakou, D. Vourloumis, D. A. Spandidos, Comparative CYP1A1 and CYP1B1 substrate and inhibitor profile of dietary flavonoids. Bioorg Med Chem 19, 2842–2849 (2011).

106. H. J. Kim, S. B. Lee, S. K. Park, H. M. Kim, Y. I. Park, M. S. Dong, Effects of hydroxyl group numbers on the B-ring of 5,7-dihydroxyflavones on the differential inhibition of human CYP 1A and CYP1B1 enzymes. Arch Pharm Res 28, 1114–1121 (2005).

107. S. J. Juliusson, J. K. Nielsen, V. Runarsdottir, I. Hansdottir, R. Sigurdardottir, E. S. Bjornsson, Lifetime alcohol intake and pattern of alcohol consumption in patients with alcohol-induced pancreatitis in comparison with patients with alcohol use disorder. Scand J Gastroenterol 53, 748–754 (2018).

108. M. H. Kim, Flavonoids inhibit VEGF/bFGF-induced angiogenesis in vitro by inhibiting the matrix-degrading proteases. J Cell Biochem 89, 529–538 (2003).

109. W. A. Schroder, T. T. Le, L. Major, S. Street, J. Gardner, E. Lambley, K. Markey, K. P. MacDonald, R. J. Fish, R. Thomas, A. Suhrbier, A physiological function of inflammation-associated SerpinB2 is regulation of adaptive immunity. J Immunol 184, 2663–2670 (2010).

110. H. H. Hsieh, Y. C. Chen, J. R. Jhan, J. J. Lin. The serine protease inhibitor serpinB2 binds and stabilizes p21 in senescent cells. J Cell Sci 130, 3272–3281 (2017).

111. N. H. Lee, A. Cho, S. R. Park, J. W. Lee, P. Sung Taek, C. H. Park, Y. H. Choi, S. Lim, M. K. Baek, D. Y. Kim, M. Jin, H. Y. Lee, I. S. Hong, SERPINB2 is a novel indicator of stem cell toxicity. Cell Death Dis 9, 724 (2018).

112. H. T. Zhang, Z. G. Zha, J. H. Cao, Z. J. Liang, H. Wu, M. T. He, X. Zang, P. Yao, J. Q. Zhang, Apigenin accelerates lipopolysaccharide induced apoptosis in mesenchymal stem cells through suppressing vitamin D receptor expression. Chin Med J (Engl) 124, 3537–3545 (2011).

113. S. Segaert, S. Courtois, M. Garmyn, H. Degreef, R. Bouillon, The flavonoid apigenin suppresses vitamin D receptor expression and vitamin D responsiveness in normal human keratinocytes. Biochem Biophys Res Commun 268, 237–241 (2000).

114. C. Boeckx, K. Op de Beeck, A. Wouters, V. Deschoolmeester, R. Limame, K. Zwaenepoel, P. Specenier, P. Pauwels, J. B. Vermorken, M. Peeters, G. Van Camp, M. Baay, F. Lardon, Overcoming cetuximab resistance in HNSCC: the role of AURKB and DUSP proteins. Cancer Lett 354, 365–377 (2014).

115. N. Al Shukor, R. Ravallec, J. Van Camp, K. Raes, G. Smagghe, Flavonoids stimulate cholecystokinin peptide secretion from the enteroendocrine STC-1 cells. Fitoterapia 113, 128–131 (2016).

116. M. A. Hasnat, M. Pervin, J. H. Lim, B. O. Lim, Apigenin Attenuates Melanoma Cell Migration by Inducing Anoikis through Integrin and Focal Adhesion Kinase Inhibition. Molecules 20, 21157–21166 (2015).

117. R. Ginwala, E. McTish, C. Raman, N. Singh, M. Nagarkatti, P. Nagarkatti, D. Sagar, P. Jain, Z. K. Khan, Apigenin, a Natural Flavonoid, Attenuates EAE Severity Through the Modulation of Dendritic Cell and Other Immune Cell Functions. J Neuroimmune Pharmacol 11, 36–47 (2016).

118. D. Arango, K. Morohashi, A. Yilmaz, K. Kuramochi, A. Parihar, B. Brahimaj, E. Grotewold, A. I. Doseff, Molecular basis for the action of a dietary flavonoid revealed by the comprehensive identification of apigenin human targets. Proc Natl Acad Sci U S A 110, E2153–2162 (2013).

119. A. Onoufriadis, A. Shoemark, M. M. Munye, C. T. James, M. Schmidts, M. Patel, E. M. Rosser, C. Bacchelli, P. L. Beales, P. J. Scambler, S. L. Hart, J. E. Danke-Roelse, J. J. Sloper, S. Hull, C. Hogg, R. D. Emes, G. Pals, A. T. Moore, E. M. Chung, Uk10K, H. M. Mitchison, Combined exome and whole-genome sequencing identifies mutations in ARMC4 as a cause of primary ciliary dyskinesia with defects in the outer dynein arm. J Med Genet 51, 61–67 (2014).

120. A. Delprato, B. Bonheur, M. P. Algeo, P. Rosay, L. Lu, R. W. Williams, W. E. Crusio, Systems genetic analysis of hippocampal neuroanatomy and spatial learning in mice. Genes Brain Behav 14, 591–606 (2015).

121. L. A. Higa, D. Banks, M. Wu, R. Kobayashi, H. Sun, H. Zhang, L2DTL/CDT2 interacts with the CUL4/DDB1 complex and PCNA and regulates CDT1 proteolysis in response to DNA damage. Cell Cycle 5, 1675–1680 (2006).

122. Y. T. Huang, J. O. Mason, D. J. Price, Lateral cortical Cdca7 expression levels are regulated by Pax6 and influence the production of intermediate progenitors. BMC Neurosci 18, 47 (2017).

123. A. D. Perdereau, K. Cailliau, E. Browaeys-Poly, A. Lescuyer, N. Carre, F. Benhamed, D. Goenaga, A. F. Burnol, Insulin-induced cell division is controlled by the adaptor Grb14 in a Chfr-dependent manner. Cell Signal 27, 798–806 (2015).

124. A. R. Collins, Measuring oxidative damage to DNA and its repair with the comet assay. Biochim Biophys Acta 1840, 794–800 (2014).

125. R. M. Deacon, Housing, husbandry and handling of rodents for behavioral experiments. Nat Protoc 1, 936–946 (2006).

