## Supplementary material for "Apigenin as a Candidate Prenatal Treatment for Trisomy 21: Effects in Human Amniocytes and the Ts1Cje Mouse Model"

I- **Supplementary Methods:**
1. **In Vitro Studies:**

***Human Amniocytes***

The amniocytes were obtained after clinically indicated prenatal karyotyping. As this was discarded material that was de-identified, patient consent was deemed unnecessary. Only fetal karyotype and sex were known. Second trimester amniocytes were prepared as described previously (19). All trisomy 21 samples used in this study have full T21 and no mosaic or partial trisomy samples were used. Samples were matched for sex and gestational age according to the following table:

**Supplementary Table 1: Karyotype and gestational age information of the human Trisomy 21 and euploid amniocytes pairs used in this study.**

| **Pair** | **Sample ID** | **Gestational Age** | **Karyotype** | **Sex** |
| --- | --- | --- | --- | --- |
| **Pair 1** | **JJ1298** | 16 + 1/7 | 46XX, 2N | **Female Pair** |
|  | **CG16-467** | 16 + 5/7 | 47XX, T21 |  |
| **Pair 2** | **JT1275** | 16 + 2/7 | 46XX, 2N | **Female Pair** |
|  | **JG16-213** | 16 + 1/7 | 47XX, T21 |  |
| **Pair 3** | **KM1170** | 19 + 5/7 | 46XX, 2N | **Female Pair** |
|  | **BB16-225** | 19 + 6/7 | 47XX, T21 |  |
| **Pair 4** | **CV1166** | 15 + 4/7 | 46XY, 2N | **Male Pair** |
|  | **SZ16-201** | 15 + 3/7 | 47XY, T21 |  |
| **Pair 5** | **ER48** | 19 + 6/7 | 46XY, 2N | **Male Pair** |
|  | **JL779** | 19 + 4/7 | 47XY, T21 |  |
| **Pair 6** | **ML181** | 18 + 3/7 | 46XY, 2N | **Male Pair** |
|  | **BB1174** | 18 + 3/7 | 47XY, T21 |  |
| **Pair 7** | **LC847** | 17 + 3/7 | 46XY, 2N | **Male Pair** |
|  | **SG01** | 18 + 4/7 | 47XY, T21 |  |

***Apigenin Optimal Dose Selection Using Cell Proliferation Assays***

Amniocytes (10^+5^ cells) from fetuses with DS and euploid controls were plated in duplicate in 24-well plate culture dishes and incubated overnight at 37°C (20% O_2_, 5% CO_2_). The following day, cells were either left untreated or treated with five different concentrations of apigenin (1, 2, 3, 4 and 5 µM). Cell culture media (AmnioMax C-100 Complete, Thermo Fisher Scientific, Waltham, MA) and apigenin-containing AmnioMax C-100 Complete media were freshly prepared and changed daily during the treatment periods.

Cell proliferation was evaluated using the CellTiter 96^®^ Aqueous cell proliferation assay according to the manufacturer’s instructions (Promega, Madison, WI). Absorbance at 490 nm was measured using the Synergy 2 well plate reader (BioTek®, Winooski, VT). Results were normalized to 100 % for untreated cells for further comparison with apigenin-treated cells. Automatic cell counts were performed using the Scepter™ 2.0 Handheld Automated Cell Counter and the 60 µM sensors (EMD Millipore, Billerica, MA).

***Oxidative Stress and Antioxidant Capacity***

Amniocytes were incubated with media alone or apigenin for three consecutive days, with media and drug prepared freshly and changed daily. At the end of the treatment, efficacy was evaluated using cell proliferation, antioxidant capacity, oxidative stress and global gene expression as the endpoints.

The Comet assay or Single Cell Gel Electrophoresis (SCGE) uses a DNA-binding fluorescent dye. Because of oxidative stress, damaged DNA extrudes from the nucleus making a “tail”. The assay calculates the relative amount of DNA in the “tail” versus the nucleus “head” (123). The Comet assay was performed using the CometAssay® kit according to the manufacturer’s instructions (Trevigen, Gaithersburg, MD).

To measure the physiological responses to oxidative stress before and after apigenin treatment, cells were centrifuged (1,100 g for 5 min), rinsed with ice cold 1X PBS and centrifuged again at 1,100 g for 5 min. The cell pellet was used for protein extraction using the NucleoSpin RNA/Protein extraction kit (Macherey-Nagel, Bethlehem, PA), and protein concentration was determined using the Pierce™ BCA protein assay kit (Thermo Fisher Scientific, Cambridge, MA). 5,000 µg of total protein was used to measure the total antioxidant capacity using the OxiSelect™ Total Antioxidant Capacity (TAC) assay kit according to the manufacturer’s instructions (CellBiolabs, San Diego, CA).

***RNA Extraction and Microarray Hybridization***

For gene expression studies, amniocytes were incubated with media alone or 2 µM apigenin for three consecutive days. Cells were then rinsed with ice cold PBS 1X (-Ca^2+^, -Mg^2+^) and centrifuges at 1000 RPM for 5 min. The cell pellet was used for RNA extraction using NucleoSpin RNA/Protein extraction (Macherey-Nagel, Bethlehem, PA). RNA was processed and hybridized on the GeneChip® Human Transcriptome HT 2.0 array according to the manufacturer’s instructions (Affymetrix, Santa Clara, CA). Each array corresponded to labeled cDNA from one amniocyte culture.

Array data were normalized using the oligo R package. Of the 70492 probe sets on the array, only the 42935 gene-coding probe sets were used in further analyses. Results were further visualized using a Principal Component Analysis (PCA) as well as heatmap combined with hierarchical clustering. For pathway analyses, the 42935 probe sets were first collapsed to 33721 unique genes to remove gene-level redundancy. After that pathway analyses were carried out using the Database of Annotation, Visualization, and Integrated Discovery (DAVID), Ingenuity Pathway Analysis (IPA) and Gene Set Enrichment Analysis (GSEA).

2. **In Vivo Studies:**

***Exploratory Behavior and Spontaneous Locomotor Activity***

Exploratory behavior and locomotor activity were assessed using the open field test as described previously (124). Briefly, the mouse was placed in an open field arena consisting of a white opaque plastic box 40 cm (L) X 40 cm (W) X 40 cm (H) divided into a central zone that measured 20 cm (L) X 20 cm (W) X 20 cm (H) and a periphery. Exploratory behavior was tracked during a 60 min unique trial using the Ethovision 10.5 animal tracking system (Noldus, Leesburg, VA). The total distance travelled (cm) in the center versus periphery as well as the average velocity (cm/s) were analyzed for treated and untreated groups. Data were collected as time bins of 10 minutes and as a total over the course of the experiment.

***Motor Coordination***

Motor coordination was investigated using the rotarod test (Med Associates, Fairfax, VT) using two different protocols (fixed speed on day 1 and accelerating speed on day 2). Prior to testing with the fixed speed protocol on day 1, each mouse was given 2 x 120 s practice sessions at 16 RPM. After practice, mice were tested at three different fixed speeds (16 RPM, 24 RPM then 32 RPM) for two 120 s trials at each speed and with an inter-trial interval of 15 min. On day 2, mice were tested in two trials under conditions of increasing difficulty in which the speed of the rotation gradually increased from 4 to 40 RPM over a 5-minute period. The time to fall was recorded in seconds and analyzed for each mouse.

***Hippocampal-Dependent Memory***

Hippocampal-dependant memory was analyzed using the fear conditioning test in a conditioning chamber with stainless-steel grid floor, equipped with an electric aversive stimulator, and house light, enclosed within a sound attenuating cubicle with exhaust fan (Med Associates, Fairfax, VT). On day one (training session), each mouse was individually placed for 5 min into the conditioning chamber and allowed to explore freely (habituate) for 180 seconds. Following exploration/habituation, two mild foot shocks (0.5 mA for 2 s) were administered at 180 s and 240 s. On day two (testing session), the mice were placed into the identical conditioning chamber for 5 min with no foot shocks. Each mouse was monitored for freezing (fear) behavior. The extent (or percent) of freezing was used as a measure of the animal’s memory and analyzed as time bins of 60 s and as a total over the course of the experiment using the Freeze View software (Med Associates, Fairfax, VT).

**Supplementary Figure** **1**: **Effects of apigenin on cell proliferation in T21 and euploid amniocytes.**

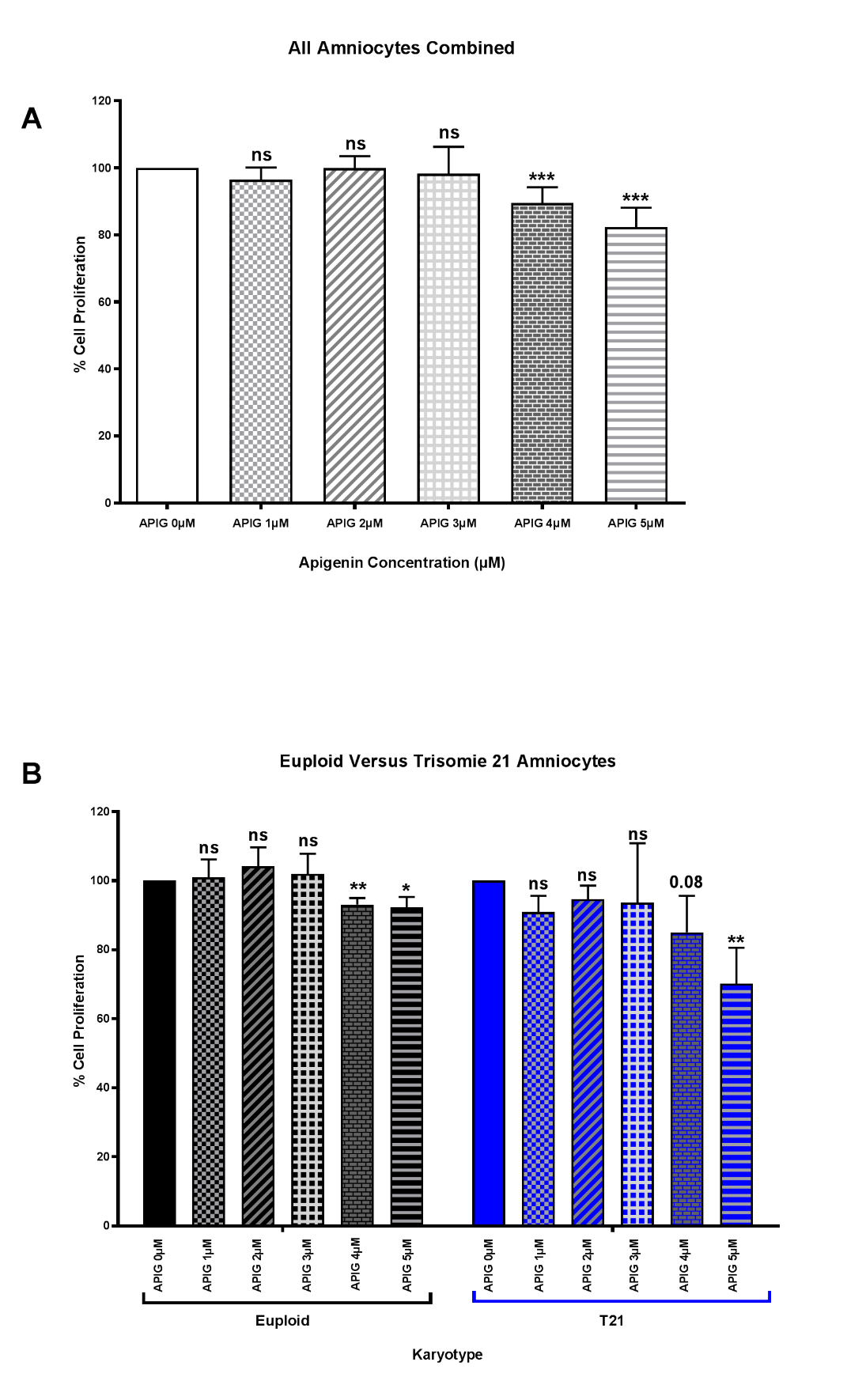

**Supplementary Figure 2:** **Effects of apigenin on natural history and growth in Ts1Cje and WT littermates.**

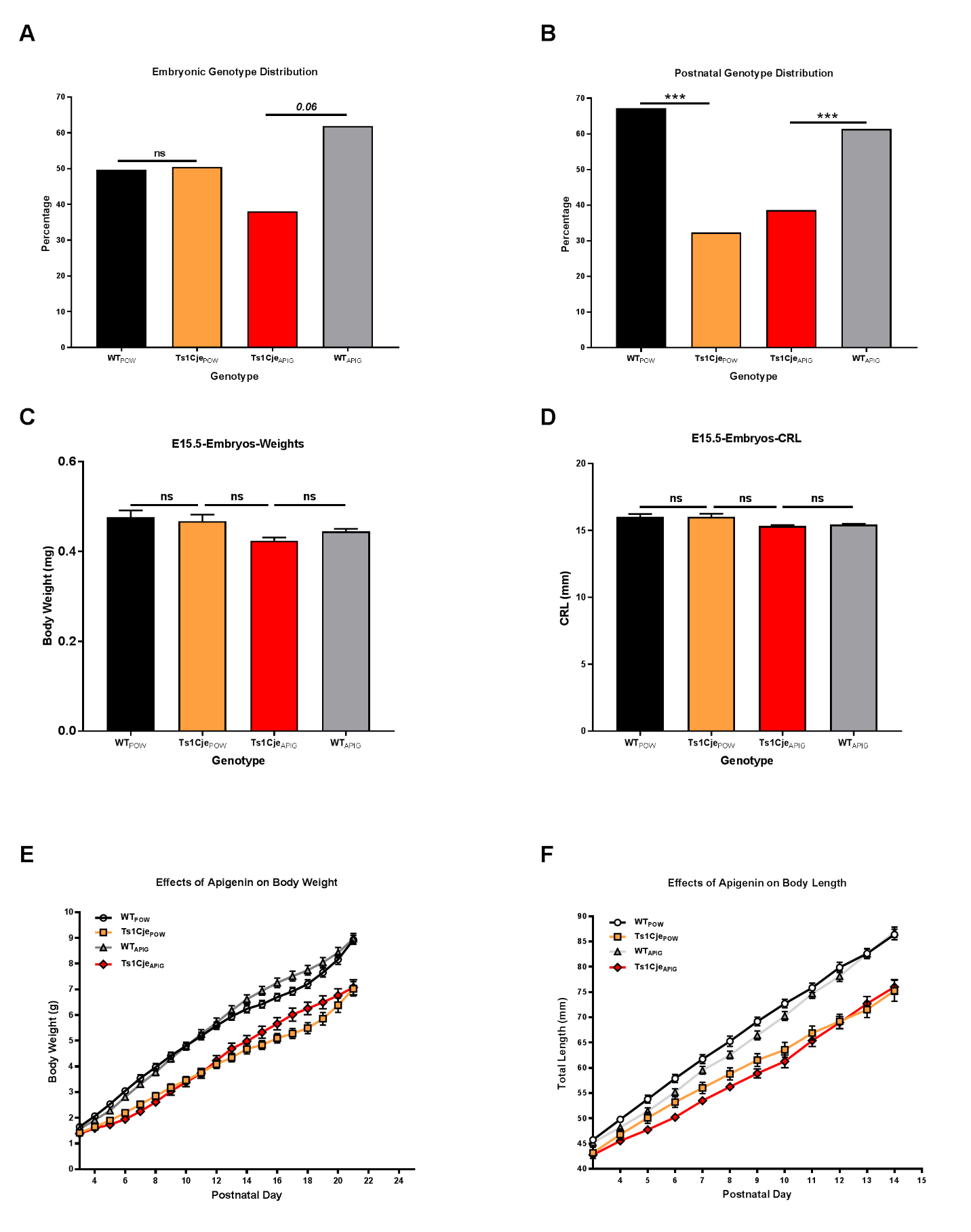

**Supplementary Figure 3: Effects of apigenin on early developmental milestones in Ts1Cje and WT littermates.**

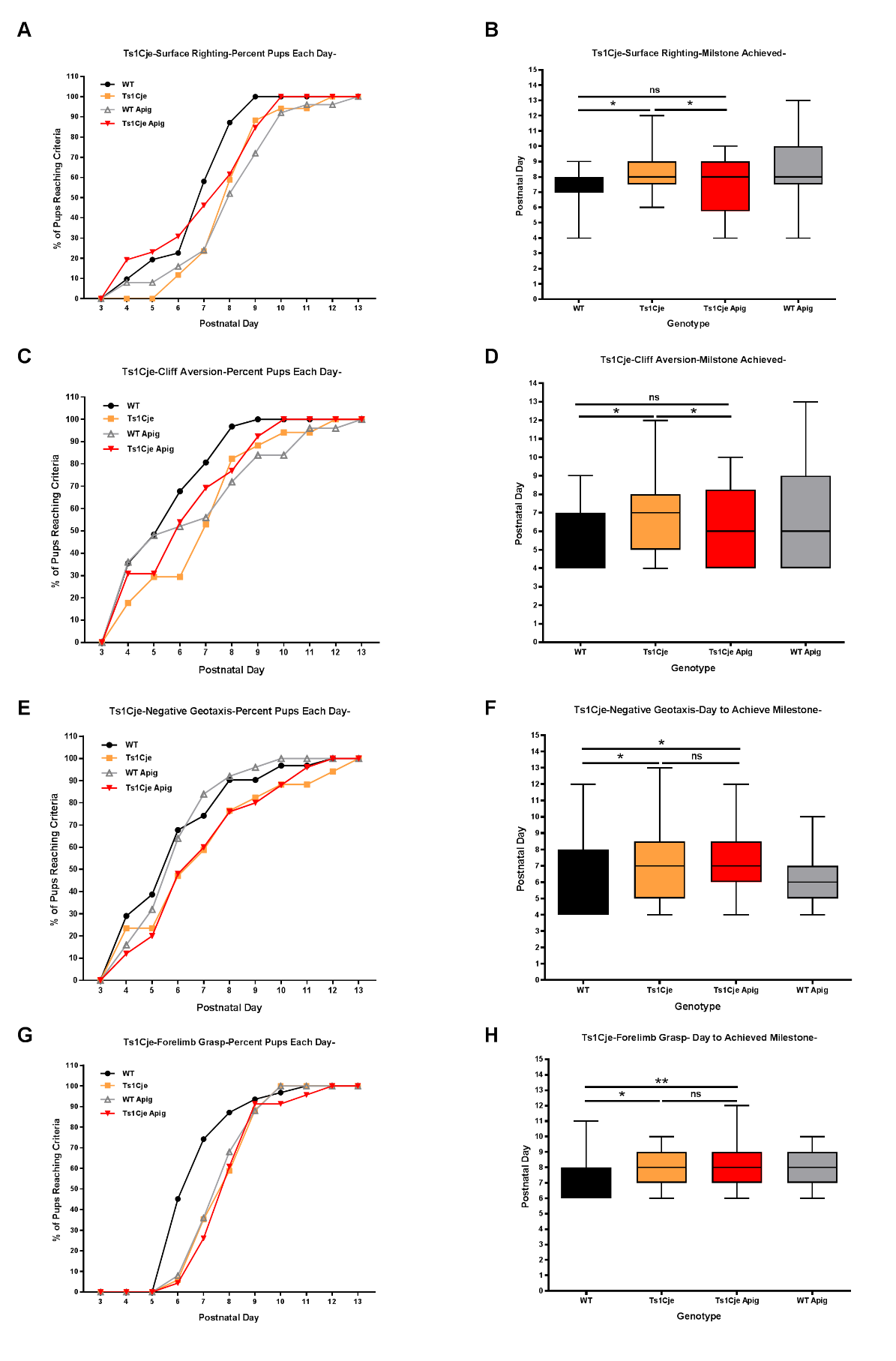

**Supplementary Figure 4: Effects of apigenin on late developmental milestones in Ts1Cje and WT littermates.**

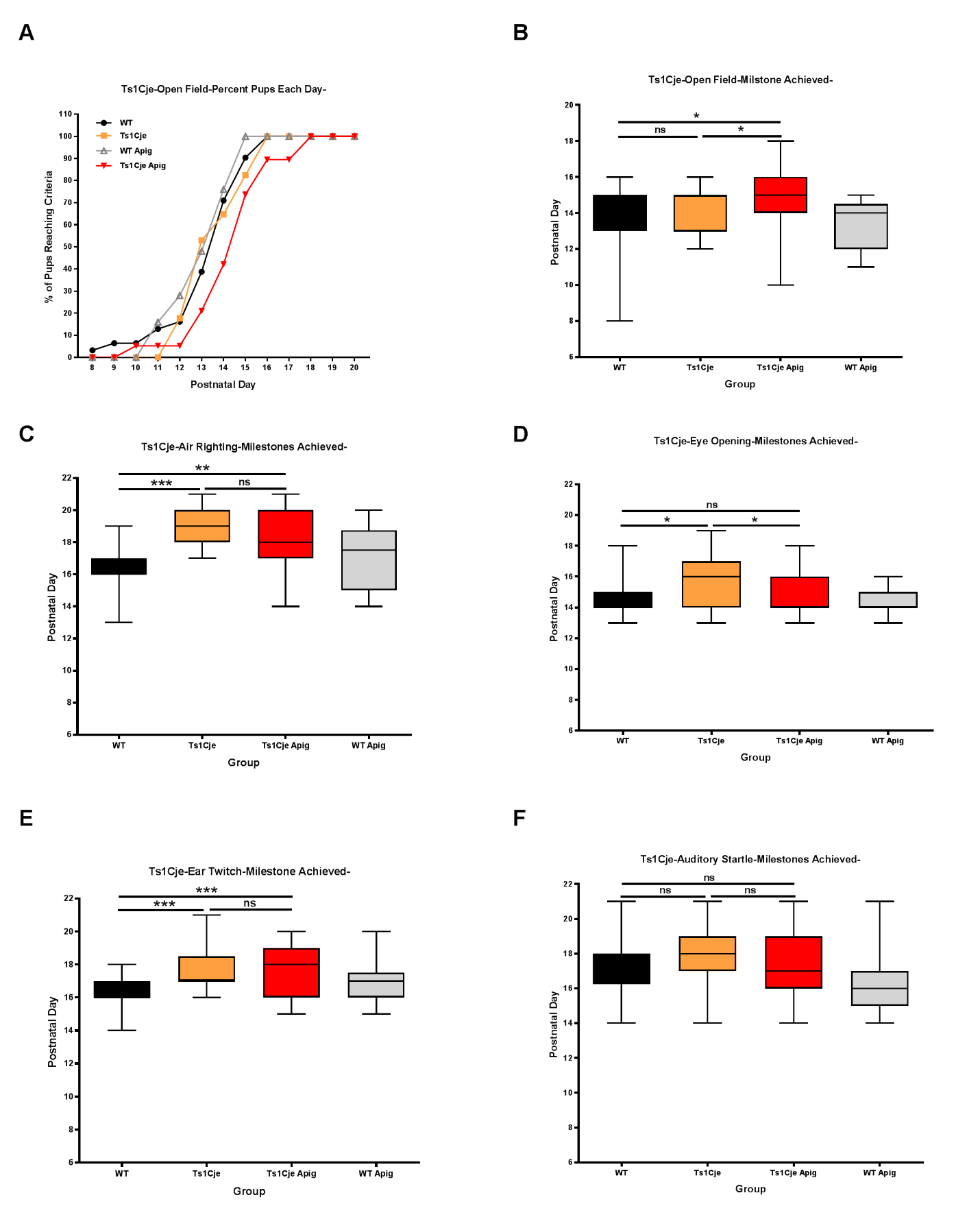

**Supplementary Figure 5**: **Adverse effects of apigenin on motor coordination in untreated and apigenin-treated adult Ts1Cje and WT males and females.**

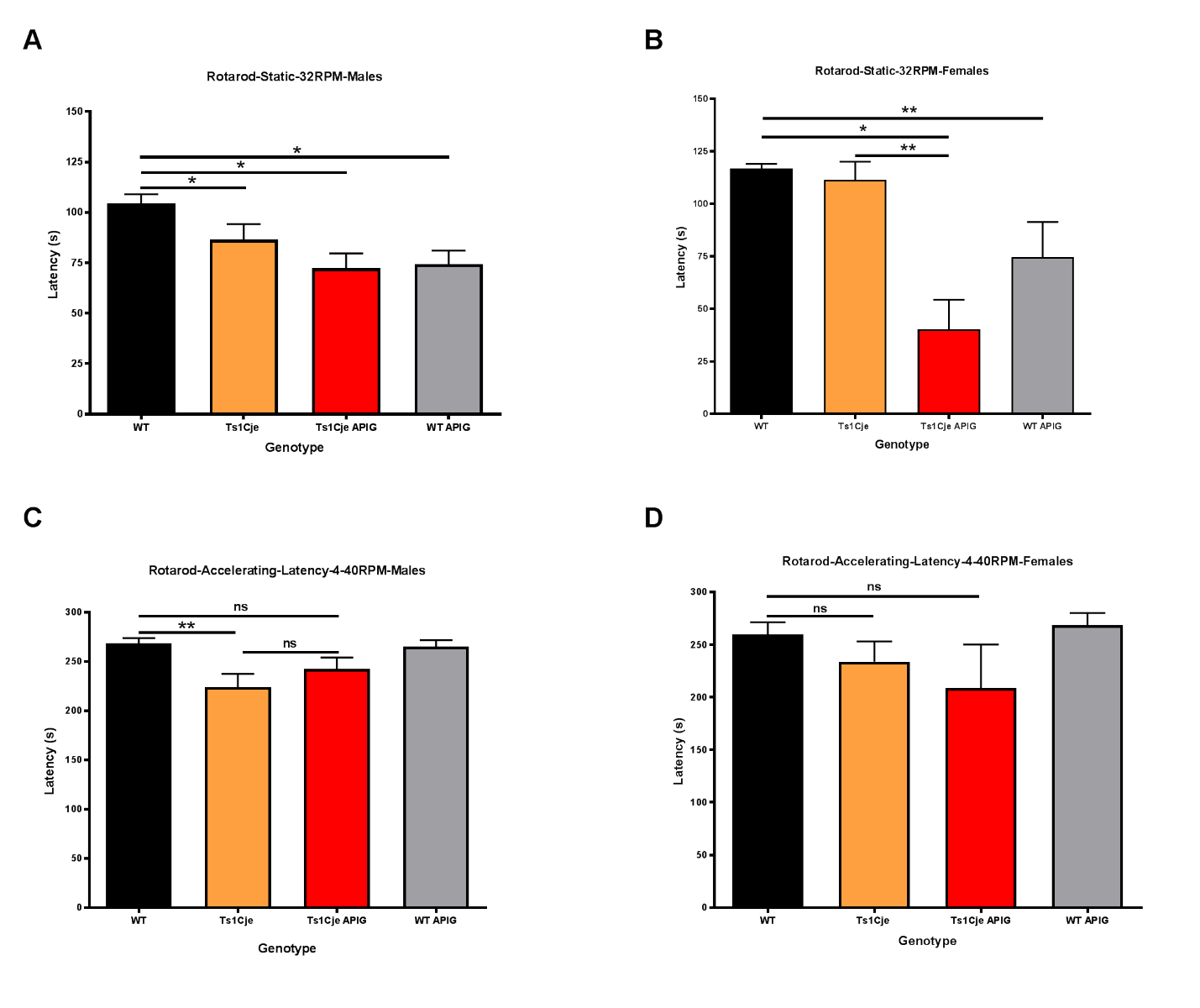

III- **Supplementary Results:**

**Supplementary Table 14: Summary of IPA Predicted Up-Stream Regulators and their Targets in Untreated and Apigenin Treated T21 Amniocytes**

| **Up-Stream Regulator** | **Signaling Pathways/ Cellular Process** | **T21 Unt/**  **Eup Untr** | | **T21 Apig/**  **T21 Untr** | | **T21 Apig/**  **Eup Untr** | | **Ts1Cje Unt/**  **Eup Untr** | | **Ts1Cje Apig/**  **Ts1Cje Untr** | | **Ts1Cje Apig/**  **Eup Untr** | |
| --- | --- | --- | --- | --- | --- | --- | --- | --- | --- | --- | --- | --- | --- |
|  |  | **Status** | **Z-Score** | **Status** | **Z-Score** | **Status** | **Z-Score** | **Status** | **Z-Score** | **Status** | **Z-Score** | **Status** | **Z-Score** |
| **VEGFA** | Angiogenesis/ Chemokine activity | Activated | 2.38 | Unchanged | 0.13 | Activated | 2.75 | Activated | 2.15 | Unchanged | 0.00 | Activated | 2.15 |
| **IFNG** | Immune Response/  Interferon Signaling | Activated | 2.02 | Inhibited | -3.09 | Partially Activated | 1.70 | Unchanged | 0.75 | Activated | 1.80 | Activated | 1.85 |
| **TNF** | Immune Response/ NFκB Signaling | Activated | 1.28 | Inhibited | -1.50 | Unchanged | 0.74 | Activation | 2.40 | Inhibited | -1.41 | Unchanged | 0.00 |
| **IRGM1** | Immune Response/  Autophagy | Unchanged | -0.18 | Inhibited | -3.59 | Inhibited | -3.60 | Unchanged | 0.00 | Inhibited | -245 | Inhibited | -2.23 |
| **TP53** | Cell stress response/  DNA Damage-Repair | Inhibited | -2.62 | Unchanged | -0.74 | Inhibited | -2.19 | Activated | 1.50 | Inhibited | -1.28 | Unchanged | -0.87 |
| **FOXM1** | Negative Regulation of G1/M transition/ DNA Damage Response | Unchanged | 0.05 | Activated | 2.7 | Activated | 2.00 | Unchanged | 0.00 | Activated | 2.01 | Activated | 2.01 |
| **FOXO1** | Insulin Signaling/Regulation of homeostasis | Unchanged | -0.51 | Activated | 3.97 | Activated | 4.35 | Unchanged | 0.00 | Activated | 2.36 | Activated | 2.17 |
| **PTGER2** | G-Protein Signaling/  Autoimmunity | Inhibited | -2.12 | Activated | 5.196 | Activated | 5.20 | Unchanged | 0.00 | Activated | 3.60 | Activated | 3.16 |
| **PTGER4** | G-Protein Signaling/  Autoimmunity | Inhibited | -1.90 | Unchanged | -0.63 | Inhibited | -1.26 | Inhibited | -1.62 | Unchanged | -0.38 | Inhibited | 0.00 |
| **HGF** | Cell proliferation/  HGF-Met Pathway | Partially Activated | 1.13 | Activated | 5.20 | Activated | 4.80 | Unchanged | 0.00 | Activated | 3.24 | Activated | 3.31 |
| **EP400** | Cell Cycle Regulation/  Positive Transcription Regulation | Unchanged | 0.25 | Activated | 3.36 | Activated | 3.36 | Unchanged | 0.00 | Activated | 3.00 | Activated | 2,45 |
| **E2F1** | Cell Cycle Regulation/ Negative Regulation of G1/S transition | Unchanged | 0.76 | Activated | 2.02 | Partially Activated | 1.87 | Unchanged | 0.00 | Activated | 2.02 | Activated | 1.71 |
| **E2F2** | Cell Cycle Regulation/ Negative Regulation of G1/S transition | Unchanged | 1.00 | Activated | 2.00 | Activated | 2.00 | Unchanged | 0.00 | Unchanged | 0.00 | Unchanged | 0.00 |
| **BNIP3L** | Pro-apoptotic Signaling/  Response to Hypoxia | Partially Inhibited | -1.29 | Inhibited | -3.36 | Inhibited | -3.87 | Unchanged | 0.00 | Inhibited | -2.98 | Inhibited | -2.24 |
| **CREB1** | RNA Polymerase II-Dependent Transcription | Unchanged | 0.80 | Unchanged | -0.88 | Unchanged | 0.99 | Unchanged | 0.00 | Activated | 2.29 | Activated | 2.30 |
| **BDNF** | Axonal Growth and Pathfinding | Unchanged | 0.00 | Unchanged | 0.00 | Unchanged | -0.08 | Unchanged | 0.00 | Inhibited | -2.00 | Inhibited | -2.36 |
| **PAX6** | Neuronal Fate Determination | Unchanged | -0.97 | Inhibited | -1.45 | Inhibited | -1.45 | Unchanged | 0.00 | Inhibited | -2.42 | Inhibited | -2.36 |
