## Supplementary figures and images for "Apigenin as a Candidate Prenatal Treatment for Trisomy 21: Effects in Human Amniocytes and the Ts1Cje Mouse Model"

### Supplementary file 2

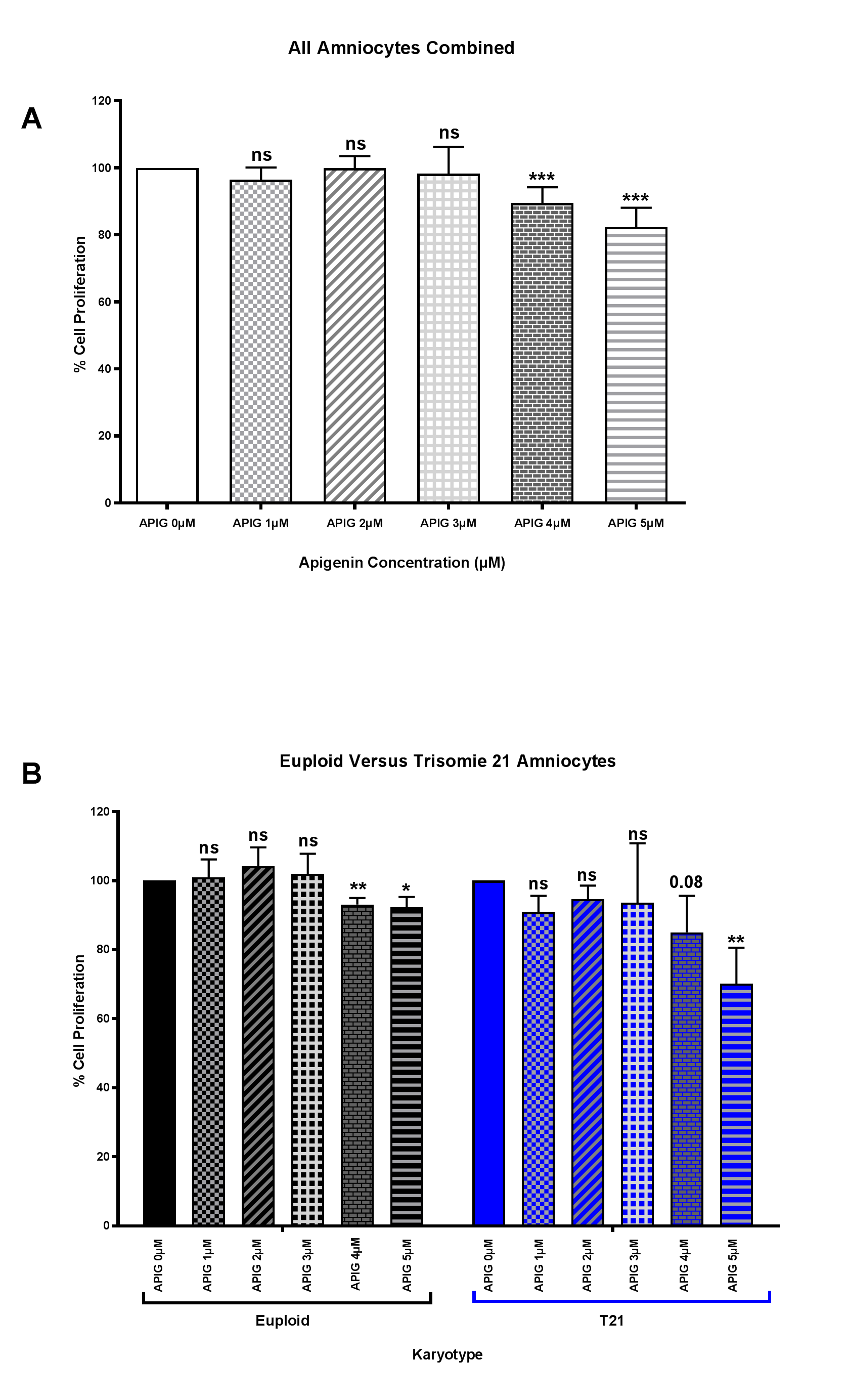

### Supplementary file 3

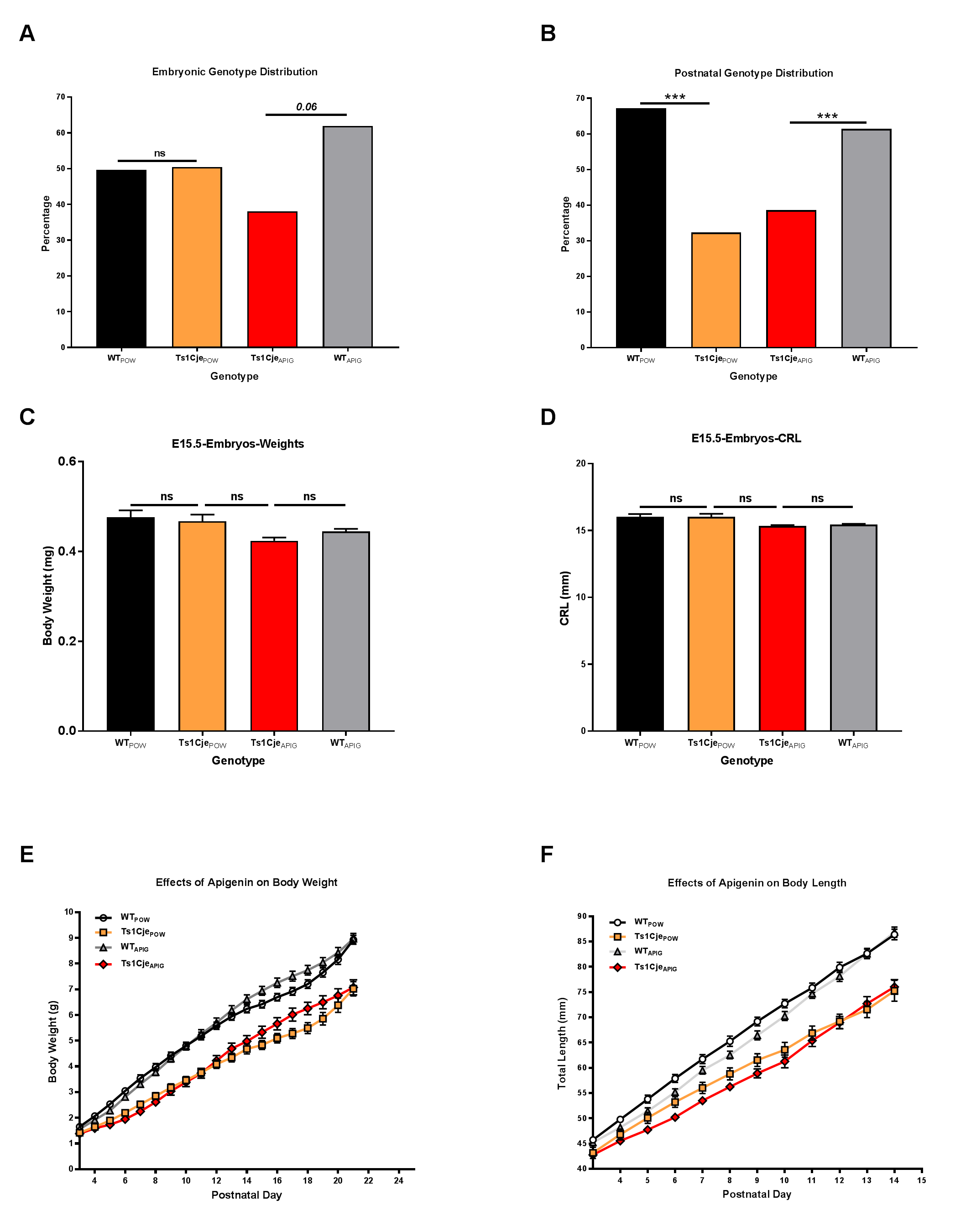

### Supplementary file 4

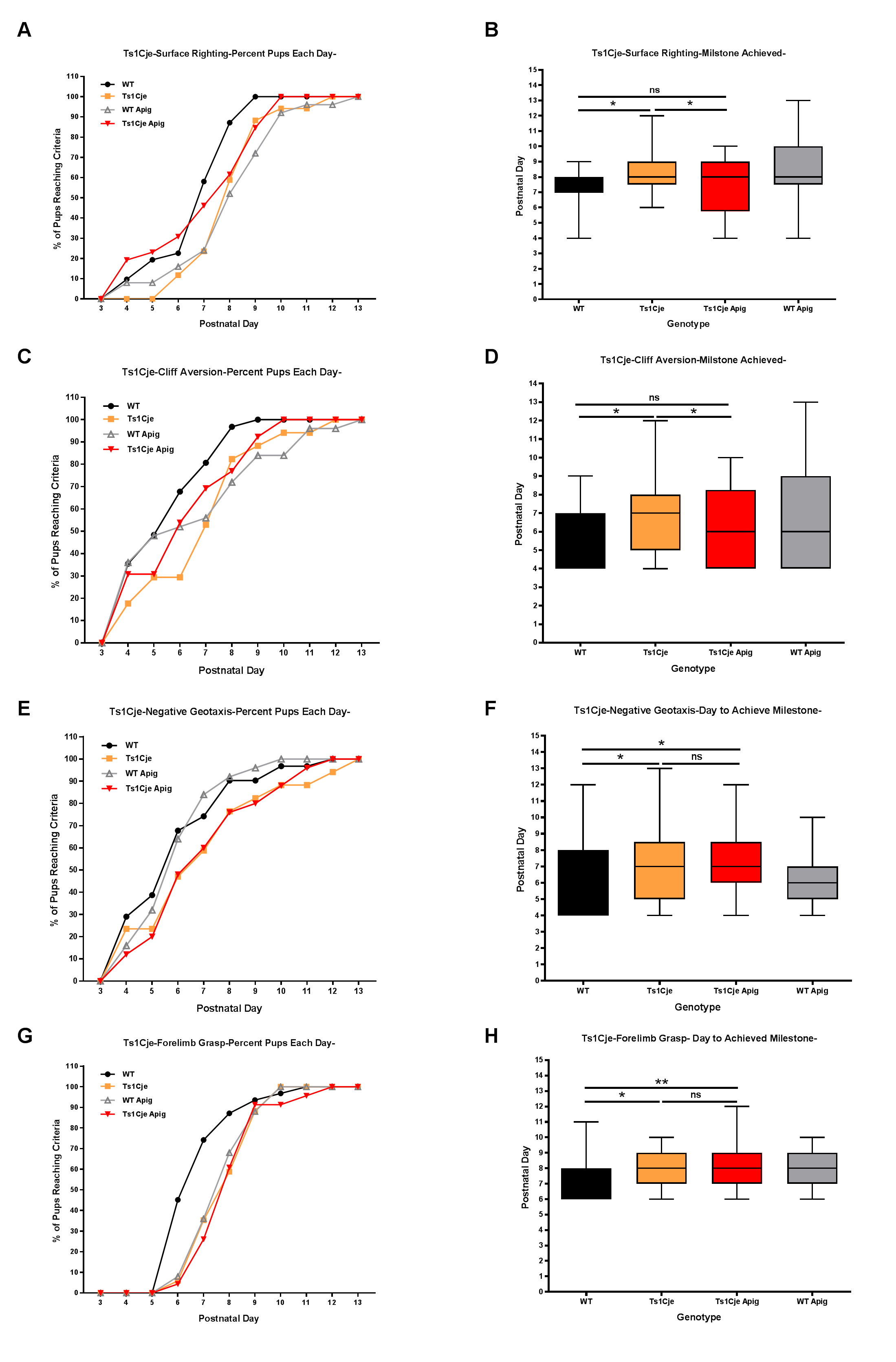

### Supplementary file 5

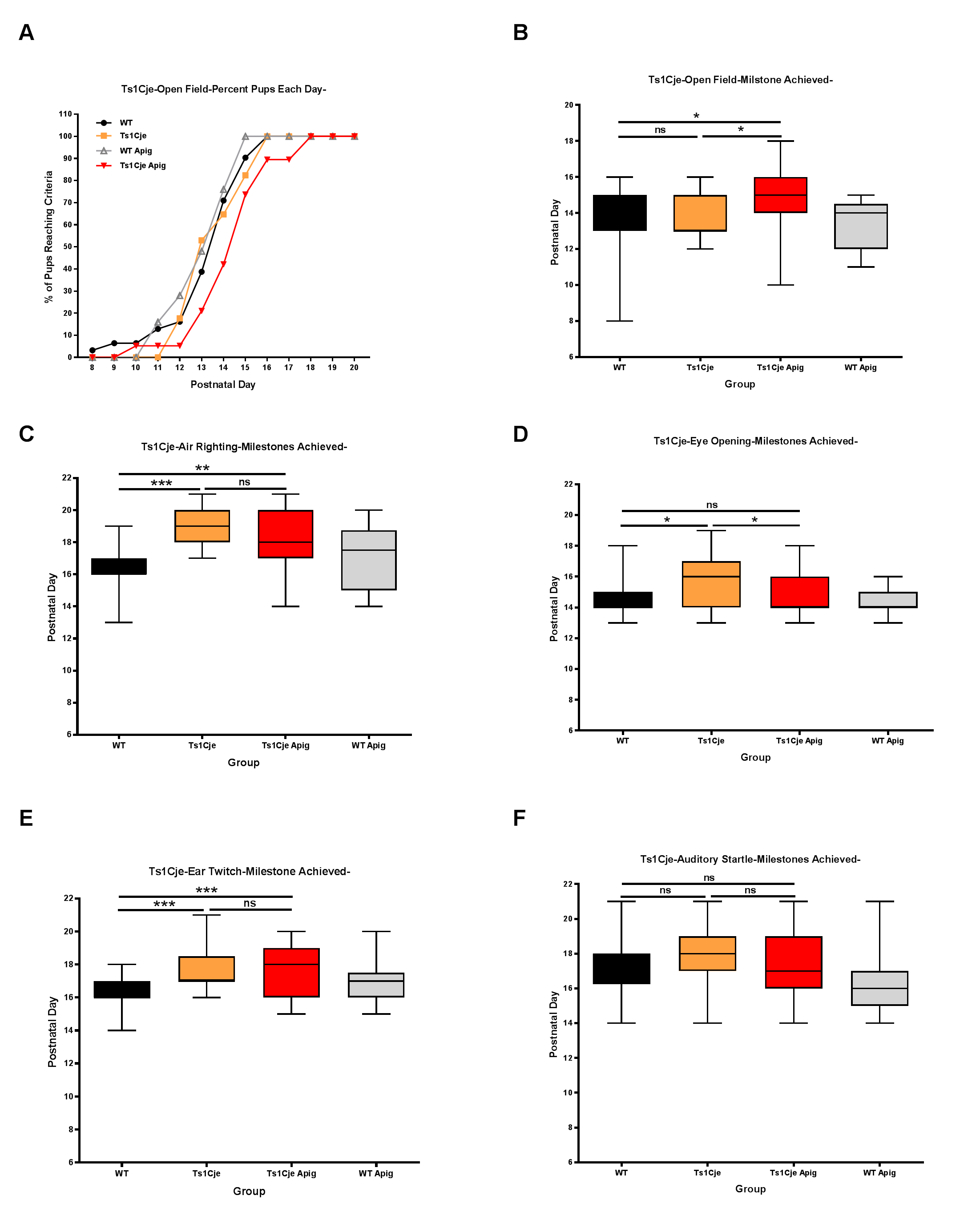

### Supplementary file 6

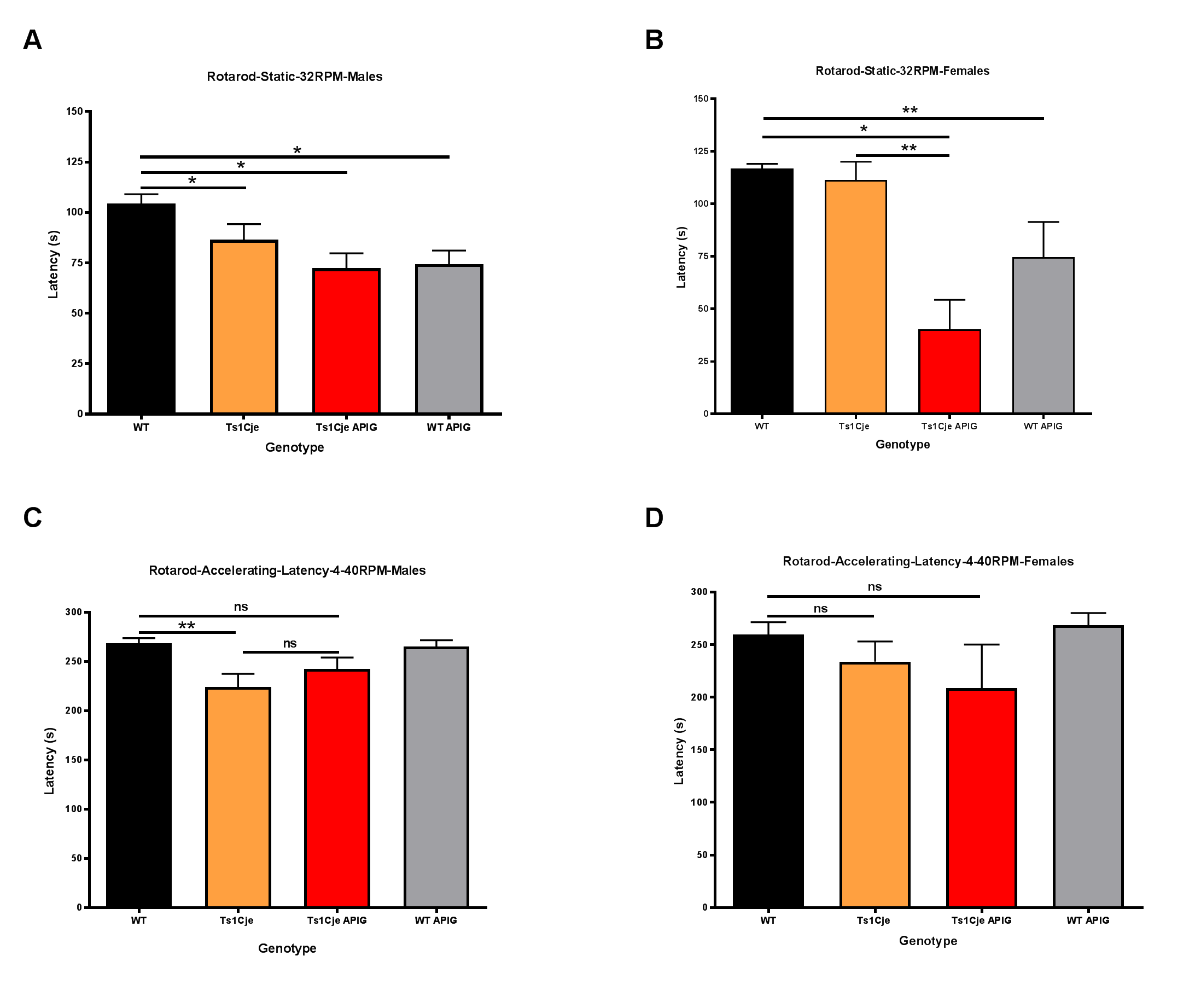
